# Dual DNA Replication Modes: Varying Fork Speeds and Initiation Rates within the spatial replication program in *Xenopus*

**DOI:** 10.1101/2024.06.21.600047

**Authors:** Diletta Ciardo, Olivier Haccard, Francesco de Carli, Olivier Hyrien, Arach Goldar, Kathrin Marheineke

## Abstract

Large vertebrate genomes duplicate by activating tens of thousands of DNA replication origins, irregularly spaced along the genome. The spatial and temporal regulation of the replication process is not yet fully understood. To investigate the DNA replication dynamics, we developed a methodology called RepliCorr, which uses the spatial correlation between replication patterns observed on stretched single-molecule DNA obtained by either DNA combing or high-throughput optical mapping. The analysis revealed two independent spatiotemporal processes that regulate the replication dynamics in the *Xenopus* model system. These mechanisms are referred to as a fast and a slow replication mode, differing by their opposite replication fork speed and rate of origin firing. We found that Polo-like kinase 1 (Plk1) depletion abolished the spatial separation of these two replication modes. In contrast, neither replication checkpoint inhibition nor Rif1 depletion affected the distribution of these replication patterns. These results suggest that Plk1 plays an essential role in the local coordination of the spatial replication program and the initiation-elongation coupling along the chromosomes in *Xenopus*, ensuring the timely completion of the S phase.

## Introduction

The faithful duplication of the genome is an essential and challenging event for all cells because it must robustly ensure efficient proliferation while maintaining genome stability. DNA replication starts from sites called replication origins; tens of thousands of origins are activated according to a regulated spatial and temporal program in each vertebrate cell to duplicate the chromosomes in a limited time window during the cell cycle. To better understand this process, the quantitative characterization of the highly heterogeneous replication dynamics constitutes an important step. Single DNA molecule data revealed that replication origins are spatially organized into clusters of two to ten that fire nearly synchronously at different times during the S phase in mammalian cultured cells and in *Xenopus* [1, 2, 3, 4, 5]. The *Xenopus in vitro* system recapitulates many aspects of cellular DNA replication. Replication-competent *Xenopus* egg extracts contain abundant maternal proteins and mimic the first rapid embryonic cell cycle when sperm nuclei are used as DNA templates [6]. We and others have shown that in this system replication initiates at 5 -15 kb intervals [7, 3, 4] and that the number of activated origins per time unit per length unit of unreplicated DNA, known as the initiation rate, increases to reach a maximum during the mid-late S phase before declining at the end of the S phase [8]. This bell-shaped curve of the initiation rate was found to be universal for the DNA replication kinetics from yeast to humans despite differences in origin specification and S phase length [9, 10]. The replication kinetics are considered to emerge from stochastic initiation in all eukaryotes [11, 12]. In *Xenopus*, we recently showed by numerical simulations that the genome could be segmented into regions of high and low probabilities of origin firing [13], similar to early and late replication timing domains in other model systems. This and other common DNA replication models assume that the replication fork speed is constant [9, 14]. However, single-molecule methods revealed that the fork speed is heterogeneous in mammalian cells [1, 15], *Xenopus* [5] and *S. cerevisiae* [16]. To ensure timely S phase completion, the fork speed and initiation rate are coordinated to compensate for varying replicon sizes or fork stalling. Several studies in mammalian cells, *Xenopus*, and budding yeast have shown that artificial fork slowing or stalling leads to the activation of dormant origins [17, 18, 19]. On the other side, decreasing initiation can lead to an increase in the fork speed [20, 21, 22, 23]. A similar fork speed and initiation rate coupling has been reported in early mouse embryos [24]. Fork speed and inter-origin distances (IODs) are very low in 2-cell embryos but increase progressively and coordinately during later developmental stages. However, the mechanism(s) underlying this correlation remains unclear. More than fifty protein factors spatially and temporally regulate the coordinated activation of replication origins. Two S-phase-specific kinases, cyclin-dependent kinases (CDK) and Dbf4-dependent kinases (DDK), are necessary at different steps for licensing and activating replication origins [25]. They are counteracted by two independent pathways to regulate the spatiotemporal program negatively. The ATR/Chk1-dependent replication checkpoint pathway inhibits the activation of late replication origins in yeasts [26, 27, 28], *Xenopus* [5, 29, 30, 31], and mammalian cells [32, 33], by targeting CDK and DDK kinases. On the other hand, the replication timing regulator Rap1-interacting factor (Rif1) has been shown to inhibit late replication in yeast [34, 35], mice [36], and human cell culture lines [37] at the level of large chromatin domains. In *Xenopus*, the depletion of Rif1 accelerates the replication program by accelerating origin cluster activation and increasing replication foci number [38]. Rif1 opposes replicative helicase activation by counteracting DDK-mediated MCM2-7 activation [39, 40, 41]. Finally, Polo-like kinase 1 (Plk1), mainly known to regulate mitosis, checkpoint recovery, and adaptation [42], is also a positive regulator of the replication program. During the S phase, Plk1 increases DNA synthesis in mammalian cells by promoting pre-replication complex loading or maintenance [43, 44]. It also promotes origin activation in the *Xenopus in vitro* system by inhibiting the replication checkpoint and Rif1 [45, 46, 47]. However, the role of these three regulatory pathways in the spatial organization of the replication process is poorly understood.

To address this question, we developed a powerful and robust analysis approach named RepliCorr, which facilitates the quantitative characterization of replication patterns measured on stretched DNA molecules during DNA combing and high-throughput optical mapping experiments. RepliCorr revealed two spatially and temporally separated replication processes in the *Xenopus in vitro* system. The first process shows a fast replication fork speed coupled with a low initiation rate, whereas the second process shows a slow replication fork speed associated with a high initiation rate. We used RepliCorr to analyze experiments in which three regulatory pathways were disrupted [31, 47, 38]. Chk1 inhibition or over-expression and Rif1 depletion did not affect the organization of these two processes. However, the depletion of Polo-like kinase 1 canceled out this dynamic separation. These results strongly suggest that Plk1 regulates the spatial replication program and the coupling between initiation and elongation in early *Xenopus* embryos to ensure the timely completion of the S phase.

## Methods and Materials

### DNA combing experiments in the Xenopus in vitro system and data analysis

DNA combing data were chosen from experiments during a control S phase and after Plk1 depletion [47], Rif1 depletion [38] or after Chk1 inhibition by UCN-01 and Chk1 overexpression [31]. Detailed experimental conditions and primary analysis are described in the respective publications. Briefly, sperm nuclei (2000 nuclei/µl) were replicated in egg extracts in the presence of biotin-dUTP naturally synchronously; genomic DNA was isolated at different times during the S phase and stretched on silanized coverslips. After immunolabelling, images were captured using a fluorescence microscope, and replication eyes were defined as the incorporation tracks of biotin-dUTP on DNA molecules. Each molecule was measured using Fiji software [48] and compiled using macros in Microsoft Excel. The replicated fraction f of each fiber was calculated as the sum of eye lengths (red tracks, Streptavidin AlexaFluor594) divided by the total DNA length (green track, anti-DNA antibody, AlexaFluor488). The initiation rate was calculated as follows:

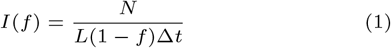

where N represents the number of new initiations defined as replication eyes smaller than 3 kb, *L* is the length of the fiber, *f* is the replicated fraction of the DNA molecule, and Δ*t* = 180*s* is the time interval in which a detectable initiation event can occur, considering that the average replication fork speed in the *Xenopus in vitro* system is ∼ 0.5*kb/min* [5]. After identifying replicated and unreplicated tracks on each DNA molecule, we constructed a binary signal where ”1” and ”0” were assigned to replicated and un-replicated units, respectively. To obtain the autocorrelation function of the fluorescence intensity profile of each molecule, we used the unbiased estimate of the cross-correlation (*xcorr*) function in Matlab (vR2013a):

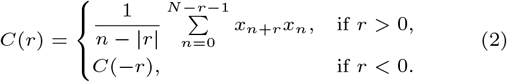

where *x*_*n*_ corresponds to the binarized signal at the position n. DNA molecules *>* 80kb were selected and ordered by the replicated fraction and grouped in bins of different sizes depending on the sample. Bins were *f*_1_ = 0 − 0.11, *f*_2_ = 0.11 −0.21, *f*_3_ = 0.21 −0.32, *f*_4_ = 0.32 −0.42, *f*_5_ = 0.42 −0.54, *f*_6_ = 0.54 − 0.64, *f*_7_ = 0.64 − 0.75. The averaged initiation rate and correlation function were calculated and plotted as a function of the averaged replicated fraction for the molecules in the bins.

### HOMARD data analysis

Images of replicating DNA molecules from sperm nuclei in egg extracts were obtained by HOMARD (High-throughput Optical Mapping of Replicating DNA) using the nanochannel array Irys® system (BionanoGenomics) as described [49] using the same fluorescent labeling strategy as in OMAR [50]. In total, 100 580 fibers from nuclei stopped in the early S phase (35 min), and 47 915 fibers from nuclei stopped in the late S phase (120 min). The fibers were visualized in blue for total DNA (Yoyo-1) and red for replicating tracks after directly incorporating AlexaFluor 647 aha-dUTP. Images were corrected for chromatic focal aberration to superimpose the blue and red channels exactly. The replicating signal detected along each stained DNA molecule was binarized using a standard thresholding method. The replication fraction *f* of each fiber was defined as the average of its binary signal. The correlation between fiber profiles was performed on the binary signal.

### Monte Carlo simulation of DNA replication process

A Monte Carlo method was used to simulate the DNA replication, as previously defined in [8]. In the simulation 100 DNA molecules were reproduced as a one-dimensional array of 150 blocks with a value 1 for replicated DNA and 0 for unreplicated DNA. Each block was considered as 1kb. At each step, the origins to activate were selected depending on the probability of initiation *P* (*t*). Initiation was allowed only in the unreplicated fraction of the simulated fibers. At each step, to reproduce DNA elongation, forks move by one block. For each simulation, a constant speed was fixed as *v* = *Nkb/min*. Then, the interval between two consecutive steps of the simulation was defined by the time necessary to replicate one block by a single fork and was set equal to 1/v. As in the KJMA models, the critical nucleus size (above which nuclei grow but below which they dissolve) is considered infinitesimal, the activation of an origin at a given position does not induce the conversion of the block value from 0 to 1. The initiation will be visible only at the following step due to the elongation.

### Model of the autocorrelation function of fluorescence profiles of replicated DNA molecules

We considered that the dynamics of DNA replication along the genome are analogous to the one-dimensional nucleation and growth process, as previously described [51, 52, 53]. The rate of origin firing per time unit and length of unreplicated DNA is temporally scale-free [53]. We then assumed this explicit form for the initiation frequency as a function of time: *I*(*t*) = *I*_0_*t*^*α*^, with *I*_0_ ≥ 0 and *α* ≥ 0. This expression is a good approximation for the increasing region of the initiation frequency, to which we restricted the analysis. By considering the work of Sekimoto [54], the replicated fraction of a molecule as a function of time t was expressed as:

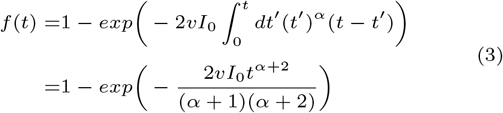

The autocorrelation function of two points separated by a distance r at a certain time t was expressed as:

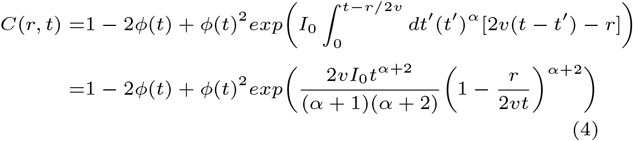

where *v* is replication fork speed and *ϕ*(*t*) = 1 − *f* (*t*) is the unreplicated fraction of a fiber. Eq. (4) is valid for *r < l*_*max*_ = 2*vt*, where *l*_*max*_ represents the maximum replication eye length present at time *t*. Calculations are detailed in Supplementary Methods.

### Parameters optimization

The replication parameters *v, I*_0_ and *α* of the model were estimated given the experimental frequency of initiation *I*(*f*)as a function of the replicated fraction *f* and the correlation function *C*(*r, f*) for different replicated fractions *f* as a function of *r*. The time was obtained from the analytical inversion of the Eq. (3) as *t* = *f*^*−*1^(*v, I*_0_, *α*). We used the genetic optimization algorithm on the Matlab platform (vR2012a) for parameter optimization. The fitness function was defined as the reduced 𝒳^2^. In the genetic algorithm, we used ten subpopulations of 10 individuals with a migration fraction of 0.1 and a migration interval of fifty steps. Each individual defined a set of variables for the fit, and the subpopulation variables were chosen within the bounds reported in Table 1. At each generation, one elite child was selected for the next generation. The rest of the population comprised 60% of children obtained after a scattered crossover between two individuals chosen by roulette wheel selection and 40% of children obtained by uniform mutation. The genetic algorithm was stopped after 3000 generations or if the fitness function attained a value of 0.5.

**Table 1.** Lower and upper bounds of the fit variables.

| Variable | Lower bound | Upper bound |
| --- | --- | --- |
| $v$ | 1e-10 | 10 |
| $I_0$ | 1e-15 | 1 |
| $\alpha$ | 0 | 5 |

### Principal component analysis and agglomerative hierarchical clustering

DNA molecules *>* 80kb in length were ordered by the replicated fraction and grouped in bins of different sizes depending on the sample. A matrix was obtained for each replicated fraction bin: each row represented the correlation function of one molecule in the bin; each column represented the value of all the correlation functions for a specific distance r. We considered r in the interval 0 to 25 kb. To reduce the dimensionality of the dataset, we performed principal component analysis on the matrix of the correlation functions with the *princomp* function in Matlab. We then used agglomerative hierarchical clustering to group the correlation functions according to different numbers of clusters and the silhouette values as clustering evaluation criteria. More precisely, the pairwise distance between pairs of correlation functions in a given bin was calculated as one minus the correlation coefficient between the two curves (*pdist* function with *correlation* distance matrix) to obtain a ‘distance vector’. An agglomerative hierarchical cluster tree was created using a weighted average distance (WPGMA) linkage method (*linkage* function). Finally, we used the *cluster* function to cut the hierarchical tree into two to five clusters. For each configuration, the clustering solution was evaluated by calculating the average silhouette values of each data point. The silhouette value for each point was obtained as follows:

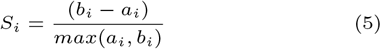

where *a*_*i*_ is the average distance from the *i*-th point to the other points in the same cluster, and *b*_*i*_ is the minimum average distance from the *i*-th point to points in a different cluster, minimized over clusters. The distance was calculated as one minus the correlation coefficient between points. The silhouette value for a point in a cluster measures how it is similar to other points in its cluster compared to points in different clusters and ranges from -1 to +1.

## Results

### Application of the auto-correlation function to replication patterns of single DNA molecules

To investigate the replication dynamics at the single molecule level, we used DNA combing data for the unperturbed S phase in the *Xenopus in vitro system* (Figure 1A) [47]. Next, the characteristics of observed replication patterns were obtained by filtering each DNA molecule using an auto-correlation function (Figure 1B), as described in the Methods. Auto-correlation filtering quantifies the spatial regularity of the observed pattern and extracts a characteristic length over which a particular signal feature holds. This widely used filtering has been successfully applied, for example, to analyze the nucleosome positioning patterns [55]. Here, we developed a new method to analyze the fluorescent signals from combed DNA molecules based on the KJMA model, which is detailed in the Methods and Supplementary Methods. We obtained an analytical expression for the autocorrelation (Eq. 4). Applying this method to analyze the replication pattern of a single DNA molecule allowed us to extract the regularity of the replicated tracks and the average correlation distance *d* over which the replication signal holds. First, to check the general feasibility of this approach, we applied the autocorrelation function to a simulated data set of replicated DNA molecules, as detailed in Methods. In this data set, we used constant values for initiation rate and fork speed and sorted the simulated replicated DNA fibers with respect to their replication fraction into seven non-overlapping bins. We then calculated each molecule’s auto-correlation profile *C*(*r, f*) as a function of the lag distances *r* and their average replication fraction, *f* (Figure 1C, blue). A good fit of the simulated data was obtained with the analytical expression of Eq. 4 (red), giving correct parameters (Supplementary Table S1). As expected by the formula, we found that at 0% replication, the *C*(*r, f*) was 0, whereas *C*(*r, f*) tended towards 1 as replication approached 100%. The maximal correlation value of each bin corresponds to the average replicated fraction of each bin. The slope for small lag distances r is given by the exponential decay constant 1/2*vt* in the expression of the correlation function (Eq. 4). With increasing time or replicated fractions, the slope becomes shallower. Indeed, as the replication progresses, the average replicated track size increases. Hence, at short lag distances (*r* ≤ 2*vt*), *C*(*r, f*) highlights processes regulating DNA elongation. At long lag distances (*r* ≥ 2*vt*), *C*(*r, f*) reflects the distribution of activated replication origins on each DNA molecule.

**Figure 1.**
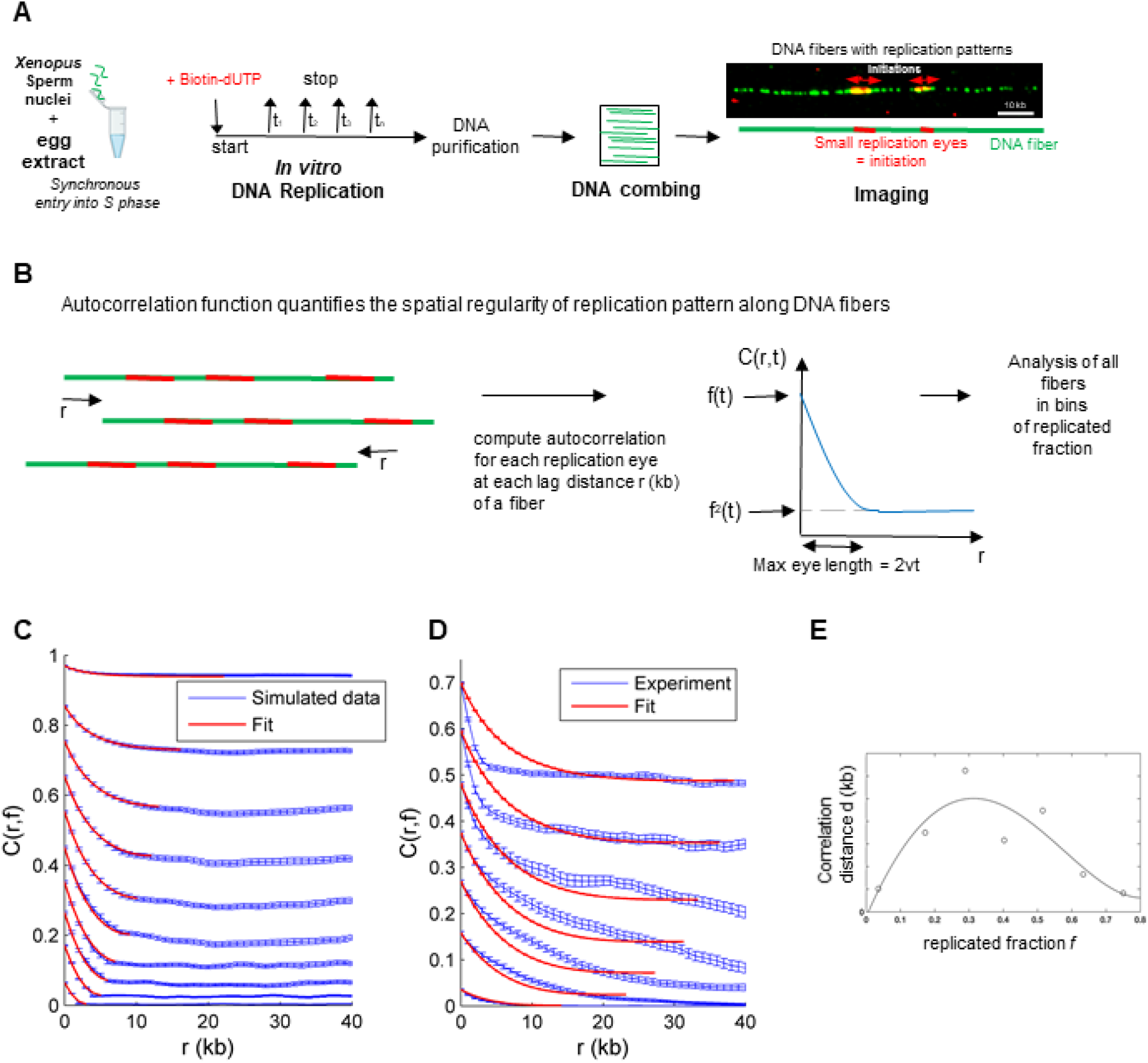
Filtering of replication patterns of single DNA molecules from DNA combing experiments during unperturbed S phase by an auto-correlation function *C*(*r, f*). (**A**) Workflow of DNA combing experiments in *the Xenopus in vitro* system. Sperm nuclei were incubated in egg extract in the presence of biotin-dUTP; replication reactions were stopped at different times during the unperturbed, naturally synchronous S phase; DNA was purified and stretched onto coverslips. Replicated tracks on single DNA fibers were revealed by fluorescence microscopy after immunolabelling (replication eyes (red), DNA molecule (green)). (**B**) Auto-correlation function C(r) measures the spatial regularity of replication patterns of a DNA molecule and its shifted copies as a function of the lag distance r. (**C**) *C*(*r, f*) profile for a simulated data set (mean with error, blue) for constant *I*(*f*) (0.03 kb^*−*1^ min^*−*1^) and constant *v* (1 kb/min) and fit (red) with one process at different bins of replicated fractions f. (**D**) Mean *C*(*r, f*) profiles (blue, with standard deviation) for three independent control DNA combing experiment at different bins of replicated fractions f and fit with one process using the KJMA model (red). (**E**) Variation of the correlation distance, d, with replication fraction. Calling *C*_*b*_(*f*), the baseline of *C*(*r, f*), d for each f is defined as *C*(*d, f*) = 1*/*2(*C*(0, *f*) *− C*_*b*_(*f*)). The open circles are experimental data, and the black curve is a 4th-order polynomial smoothing of data used to guide eyes.

We then applied this method to experimental data from three independent DNA combing experiments from the unperturbed S phase. We sorted the experimental replicated DNA molecules with respect to their replicated fraction into seven non-overlapping bins (see Methods for values). We then calculated for each DNA molecule the auto-correlation profile *C*(*r, f*) as a function of the lag distances *r* and their average replication fraction, *f*, (Figure 1D) and compared the experimental *C*(*r, f*) profiles to the simulated *C*(*r, f*) profiles (Figure 1C). As expected, as the replication degree of each bin increased, the slope of the averaged auto-correlation function *C*(*r, f*) became shallower for *r* ≤ 2*vt*. However, after the replication reached 40%, the slope of *C*(*r, f*) became sharper in the experimental profiles, whereas the slope in the simulated profiles became flatter. In addition, the fit of the experimental data with the correlation expression (red) did not reproduce the experimental profiles (blue), as it did in Figure 1C. To better visualize the slope change, we calculated a correlation distance d for the experimental data, showing a decrease at around 40% replication (Figure 1E). We interpret this transition as a change in the process regulating replication during the S phase. Altogether, these results suggest that a single process may not regulate the replication process but that two or more independent processes may be necessary to explain the experimental profiles.

### The replication process results from a combination of two spatially separated fast and slow processes

To investigate this transition further, it is necessary to understand how to link the auto-correlation profiles of each fiber to the replication process that generates the observed replication patterns. To this aim, we assumed that our set of replicated molecules contains all the patterns the replication process can produce at a given replication fraction. Next, we calculated the correlation coefficient matrices between auto-correlation profiles for each replication bin to investigate the distribution of single-molecule auto-correlation profiles along the normal S phase (Figure 2A). Each row and column represents the similarity *s* = 1−pair-wise correlation coefficient between the auto-correlation profile of one molecule and other molecules in the bin. These matrices of *s* confirm that the replication patterns are heterogeneous as scores vary [47]. Further, a closer inspection of the similarity matrices shows that their texture is granular, and lines of colors correspond to subgroups of DNA molecules with high *s* scores. This observation suggests that molecules can be clustered into subgroups of similar auto-correlation profiles. To determine how molecules should be grouped, we reduced the dimension of the *s* matrix using a Principal Component Analysis (PCA). PCA revealed that only two independent linear combinations between scores were enough to describe more than 85% of the observed variability in measured *s* scores (Figure 2B, Supplementary Figure S1). As the replication pattern of a molecule is produced by a stochastic process [11], using Kosambi-Karhunen-Lo`eve theorem [56], we concluded that only two independent stochastic processes are enough to describe the diversity of observed auto-correlation profiles. Next, we compared the averaged *C*(*r, f*) of the two independent clusters at different replicated fractions, *f*. Interestingly, while for 0 ≤ *f* ≤ 0.5, molecules with long correlation distances predominate in the population, an inversion occurred for 0.5 ≤ *f* ≤ 0.6 when molecules with short correlation distances became predominant (Figure 2C, Supplementary Figure S2). We conclude that the replication dynamics can be represented as the superposition of two independent stochastic processes. While these processes act concomitantly during the S phase, their individual effects on the replication pattern switch: we found that at low *f*, the process producing long correlation distances predominates over the process producing short correlation distances. This tendency is reversed at higher replicated fractions. Next, following the KJMA framework [52], we constructed a tractable mathematical model where the replication of a locus is induced by the simultaneous action of two independent processes (for more details, see Supplementary Methods). Each process is characterized by two parameters: the replication fork speed *v* (kb *min*^*−*1^) and the rate of replication origin activation *I*(*t*) = *I*_0_*t*^*α*^ (kb^*−*1^ min^*−*1^) per unit time per length of unreplicated DNA. Therefore, each process is characterized by three parameters *v, I*_0_, *α*. Next, using this model, we expressed the auto-correlation profile *C*(*r, f*) of a molecule with a degree of replication *f* as *C*(*r, f*) = Θ(*f*)*C*_1_(*r, f*)+(1−Θ(*f*))\**C*_2_(*r, f*), where Θ(*f*) is the mixing parameter between process 1 and 2 (0 ≤ Θ(*f*) ≤ 1). If they act alone, processes one and two create correlation profiles *C*_1_(*r, f*) and *C*_2_(*r, f*), respectively. After sorting fibers according to *f* and distributing them into seven bins, we modeled the averaged auto-correlation profile of each bin using the calculated *C*(*r, f*) (Figure 3A). For replicated fractions ≤ 0.4, the experimental correlation profiles were well reproduced by process 1 (green curve). In contrast, the average *C*(*r, f*) was predicted by process 2 (black curve) at higher replicated fractions. Fork velocity (*v*) and the initiation rate changed in opposite ways (Supplementary Table S2): process 1, named the ”fast” process, had a fast *v* and a low initiation rate, and conversely, process 2, called ”slow”, had a slow *v* and high initiation rate. More precisely, the slow process presents a nearly 6-fold lower fork speed but an 8-fold higher initiation strength *I*_0_ than the fast process. To quantitatively model *C*(*r, f*), we introduced the mixing parameter Θ(*f*) (inset in Figure 3A) that acts as an external clock regulating the transition between the fast and slow process during the S phase. This suggests that an unknown replication-independent switch triggers the change of the replication dynamics in the cell. To further control for the necessity of two independent processes with different parameters, we also fitted the auto-correlation profiles and I(*f*) from the experimental data by considering two processes with either two different fork speeds and the same initiation rate or the same fork speed and two different initiation rates (Supplementary Figure S3 A, B). In the first case, we could not fit very well the initiation rate and the time to replicate was too long to be compatible with the S phase length in this experimental system; in the second case, we could fit the initiation rate, but the time needed to replicate the fibers was again too high (Supplementary Table S2). Next, to investigate the distribution of DNA molecules between the two processes, we calculated the correlation coefficient, *ρ*, between the auto-correlation profile of each molecule *C*_1_(*r, f*) and *C*_2_(*r, f*) at a given replicated fraction. We defined the similarity distance as *s* = 1−*ρ*. To visualize this distribution, we reported the similarity distance values for each fiber to the slow and fast process on a two-dimensional graph where the x-axis represents the distance from the fast process, and the y-axis represents the distance from the slow process (Figure 3B). Since small s values indicate similarity to each process, points closer to the y-axis represent fibers from the fast process, and points closer to the x-axis represent points from the slow process. Interestingly, the data points are distributed vertically or horizontally, with few points around the diagonal. This suggests that the replication pattern of the majority of molecules is described exclusively either by the fast or the slow process alone. As the correlation distance is proportional to the fork velocity (see Supplementary methods), the fast process produces longer replicated tracks than the slow process for the same S phase length (Figure 3C). Thus, this single-molecule analysis method, which we call RepliCorr, unveils the spatial heterogeneity of fork speed and initiation rate along the genome.

**Figure 2.**
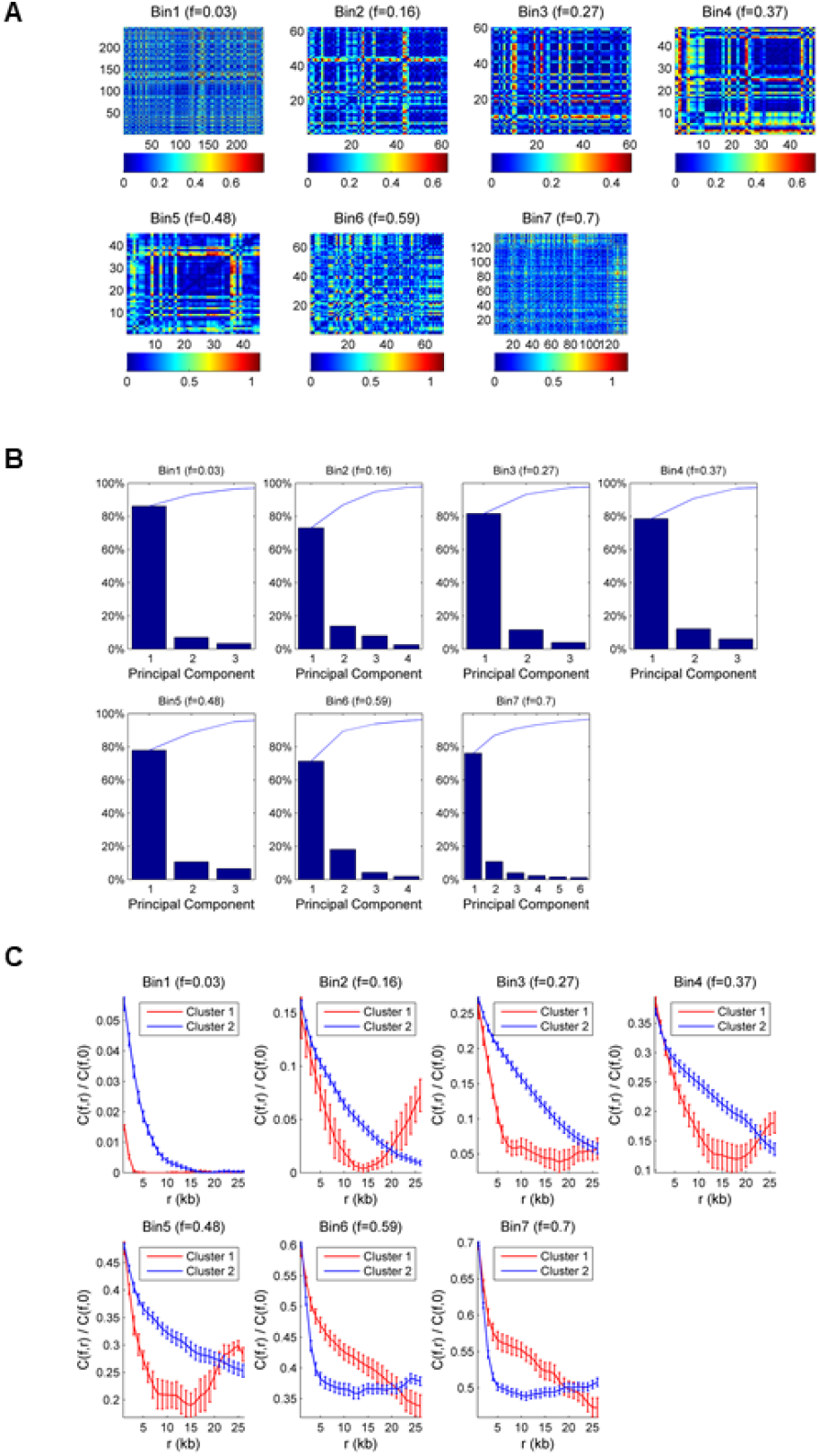
Hierarchical classification of DNA molecule’s replication patterns. (**A**) Similarity matrix between molecule’s C(r) for different replication fractions f. The similarity is measured as the *s* = 1 *−* pair-wise correlation coefficient. The color map represents the values of s for two perfectly similar C(r), s=0 (black), and for two completely different *C*(*r*), (dark red). (**B**) Pareto chart of the percent variability explained by each principal component. Each chart corresponds to a different interval of replicated fractions from 0 to 75%; the average replicated fraction is reported on top. In each chart, the bars represent the percentage of variance described by the relative principal component in descending order. The blue line represents the cumulative total. © Mean *C*(*r, f*) profiles (with standard deviation) for molecules are hierarchically classified into two similarity classes. The *C*(*r, f*) of the category containing the smaller number of molecules is in red, and the *C*(*r, f*) of the category containing the larger number of molecules is in blue. Slow decaying *C*(*r, f*) corresponds to the category with a larger correlation distance. Error bars are standard deviations.

**Figure 3.**
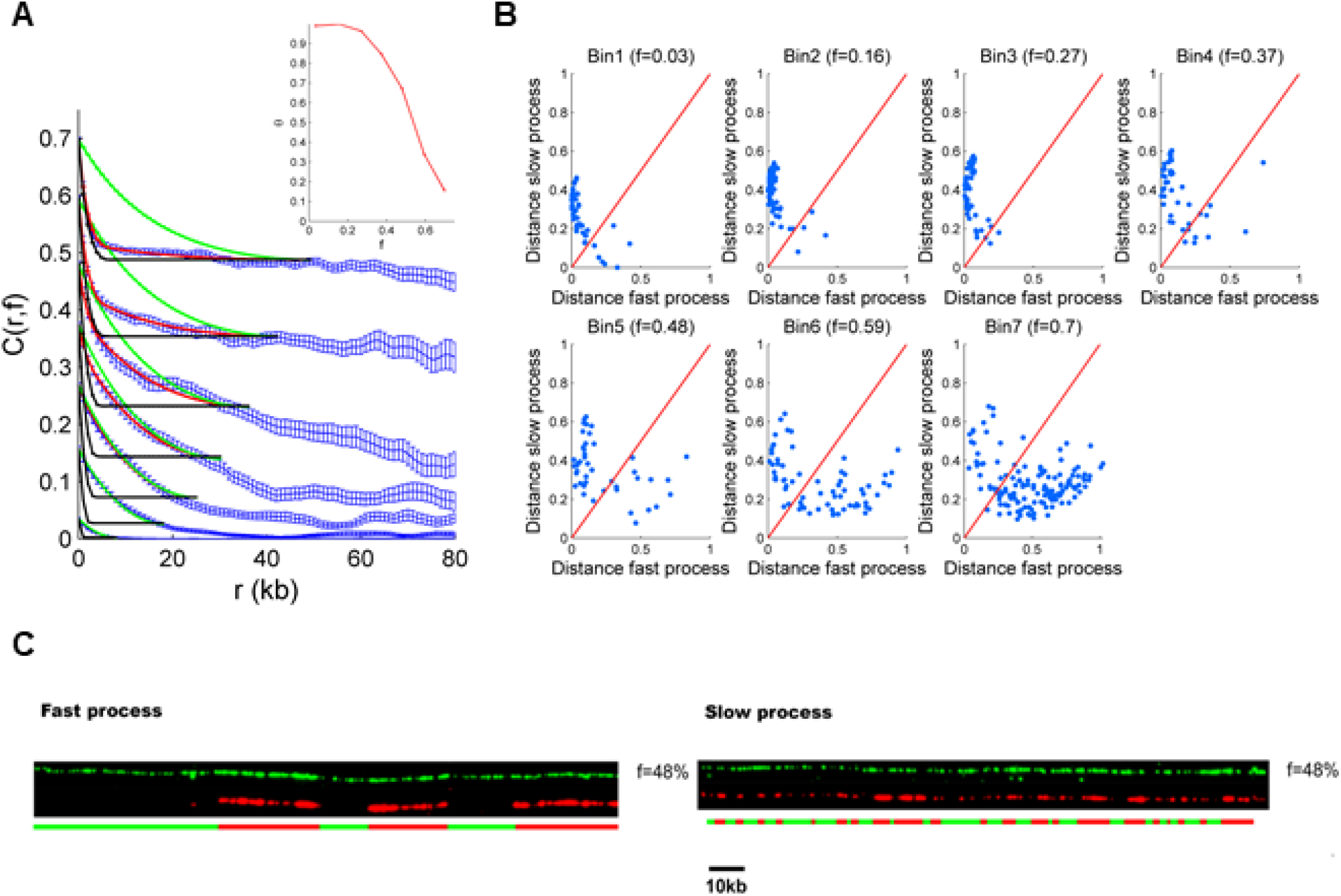
Categorization of DNA molecule’s replication patterns into two dynamical processes by RepliCorr. (**A**) Fit (red curve) of mean *C*(*r, f*) profile (blue curve, with standard deviation from three independent experiments) for each replication fraction bin calculated as *C*(*r, f*) = Θ(*f*)*C*1(*r, f*) + (1 *−* Θ(*f*)) *\* C*2(*r, f*). The green curve is the correlation profile produced by the fast fork process (C1(r,f), model 1), and the black curve is the correlation profile produced by the slow fork process (*C*2(*r, f*), model 2). The error bars are standard deviations. The inset is the Θ(*f*) profile. (**B**) Normalized correlation coefficients (*ρ*_1_, *ρ*_2_) between the molecule’s *C*(*r, f*) and *C*1(*r, f*) and *C*2(*r, f*) were calculated. The similarity distance between the molecule and each process was defined as 1-*ρ*_1_ for the fast process 1 and 1-*ρ*_2_ for the slow process and represented on a two orthogonal axis plot. The red diagonal represents points of equal similarity to the two processes. Points above the diagonal are more similar to the fast process and points below are more similar to the slow process. (**C**) Sample of replicating pattern in the fast and slow process in the same replicated fraction f=0.48. The red signal is the replicative signal, and the green is the underlying DNA molecule.

To verify whether the low-throughput of the DNA combing experiments (about 1000 fibers per condition and experiment) was sufficient for RepliCorr analysis, we analyzed unpublished high-throughput data (150 000 fibers), obtained by optical mapping of replicating *Xenopus* sperm DNA (HOMARD) in the *Xenopus in vitro* system [50] (Figure 4A, B). As for combed molecules, we modeled the auto-correlation profile of each molecule using *C*(*r, f*) = Θ(*f*)*C*_1_(*r, f*) + (1 − Θ(*f*)) * *C*_2_(*r, f*). We calculated the distance between the experimentally defined auto-correlation profile of the fiber and the slow and fast process (Figure 4C). Due to the high-throughput of the optical mapping experiment, we could apply RepliCorr analysis to the early (35 min) and the late (120 min) time points separately. As observed with the DNA combing, the data points were distributed vertically or horizontally, with few points around the diagonal direction for the early time point (blue points). However, late S phase patterns are distributed only along the axis of the slow process. This suggests that while both slow and fast processes coexisted in distinct genome regions in the early S phase, the slow process was nearly exclusive in the late S phase. Therefore, the low-throughput in DNA combing experiments neither influenced the outcome of the RepliCorr analysis nor the distribution of similarity distances along the graph representing the slow and fast processes. In addition, the high-throughput of the HOMARD analysis allowed us to visualize the temporal separation of these processes along the S phase. We conclude that DNA combing and HOMARD experiments show a clear spatial separation between the fast and the slow processes, as fibers are not distributed on the diagonal.

**Figure 4.**
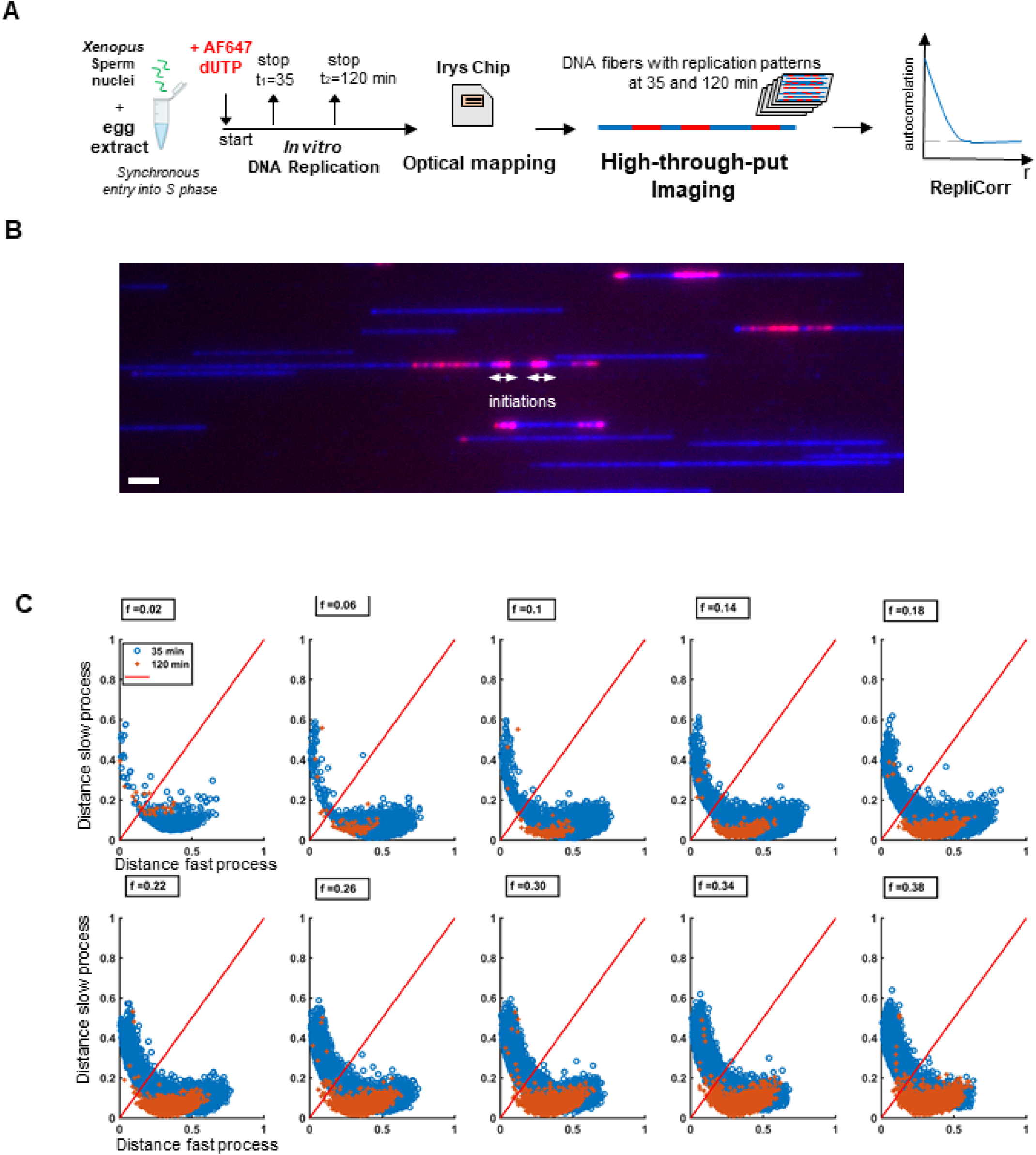
Replicorr analysis of the high-throughput optical mapping replication (HOMARD) experiments in the *Xenopus in vitro* system. (**A**) Workflow of the experiment using HOMARD: Sperm nuclei were incubated in egg extracts in the presence of AF647-aha-dUTP, stopped in early (35 min) and late S phase (120 min), DNA was isolated and separated in Irys system with Yoyo-1 stain. (**B**) Example field of view of DNA fibers with Bionano Irys system, blue, Yoyo-1, whole DNA stain, small, red replication tracks (=initiations) labeled directly by AF647-aha dUTP, early S phase (35 min), size bar 20 kb. (**C**) Normalized correlation coefficients (*ρ*_1_, *ρ*_2_) between the fibers’ C(r,f) and C1(r,f) and C2(r,f) were calculated. The similarity distance between the fiber and each process was defined as 1-*ρ*_1_ for the fast process and 1-*ρ*_2_ for the slow process and represented on a two-orthogonal axis plot. The red diagonal represents points of equal similarity to the two processes; blue points represent the early S phase (35 min), and orange points represent the late S phase (120 min).

### Depletion of Polo-like kinase 1 reduces the spatial heterogeneity of the replication profiles

To identify molecular determinants involved in the spatial separation between the fast and slow process during the S phase, we used RepliCorr to analyze DNA combing data after inhibition or depletion of different known regulators of DNA replication (Figure 5A, Supplementary Figure S4). The ATR-Chk1 dependent intra-S checkpoint pathway inhibits origin firing at the level of replication clusters in *Xenopus* [5, 31]. Inhibition of the checkpoint effector kinase Chk1 by UCN-01 or Chk1 over-expression did not alter the partition of replicating DNA molecules into two separate classes (Figure 5B, C, Supplementary Figure S5 A-B). Another important negative regulator of the replication program in *Xenopus* is Rif1 [38]. However, using RepliCorr, we found that after Rif1 depletion, the separation of molecules into the two classes was maintained (Figure 5D, Supplementary Figure S5C). Still, slightly more data points were found around the diagonal, especially in higher replicated fraction bins compared to checkpoint-inhibited conditions. This suggests that the spatial heterogeneity of patterns is only slightly reduced after Rif1 depletion. We recently found that depletion of Polo-like kinase 1 inhibited DNA synthesis via inhibition of origin activation during normal S phase in *Xenopus*, whereas the add-back of recombinant Plk1 rescued DNA replication [46, 47]. Applying RepliCorr to replication patterns after Plk1 depletion showed a dramatic change in the replication pattern partition (Figure 5E), compared to the control (Figure 3B). The long and short replicated tracks were no longer spatially and temporarily separated but coexisted on molecules with a high degree of replication. Interestingly, from the fits of the two independent processes of the correlation function and their individual parameter values (Supplementary Figure S6 A, Supplementary Table S3), we noticed that after Plk1 depletion, the parameters of the slow process tended toward those of the fast process, thus resulting in a spatially more homogeneous replication process. The rate of origin firing decreased for the slow but not the fast process, suggesting that Plk1 mainly promotes origin activation in the genomic regions governed by the slow process. Chk1 inhibition or Rif1 depletion also exclusively affected the initiation rate of the slow process but to a much lesser extent than Plk1 depletion. We conclude that Plk1 depletion has the strongest effect on separating the two replication processes highlighted by RepliCorr analysis and that Plk1 regulates the spatial organization of origin firing and fork progression along the genome.

**Figure 5.**
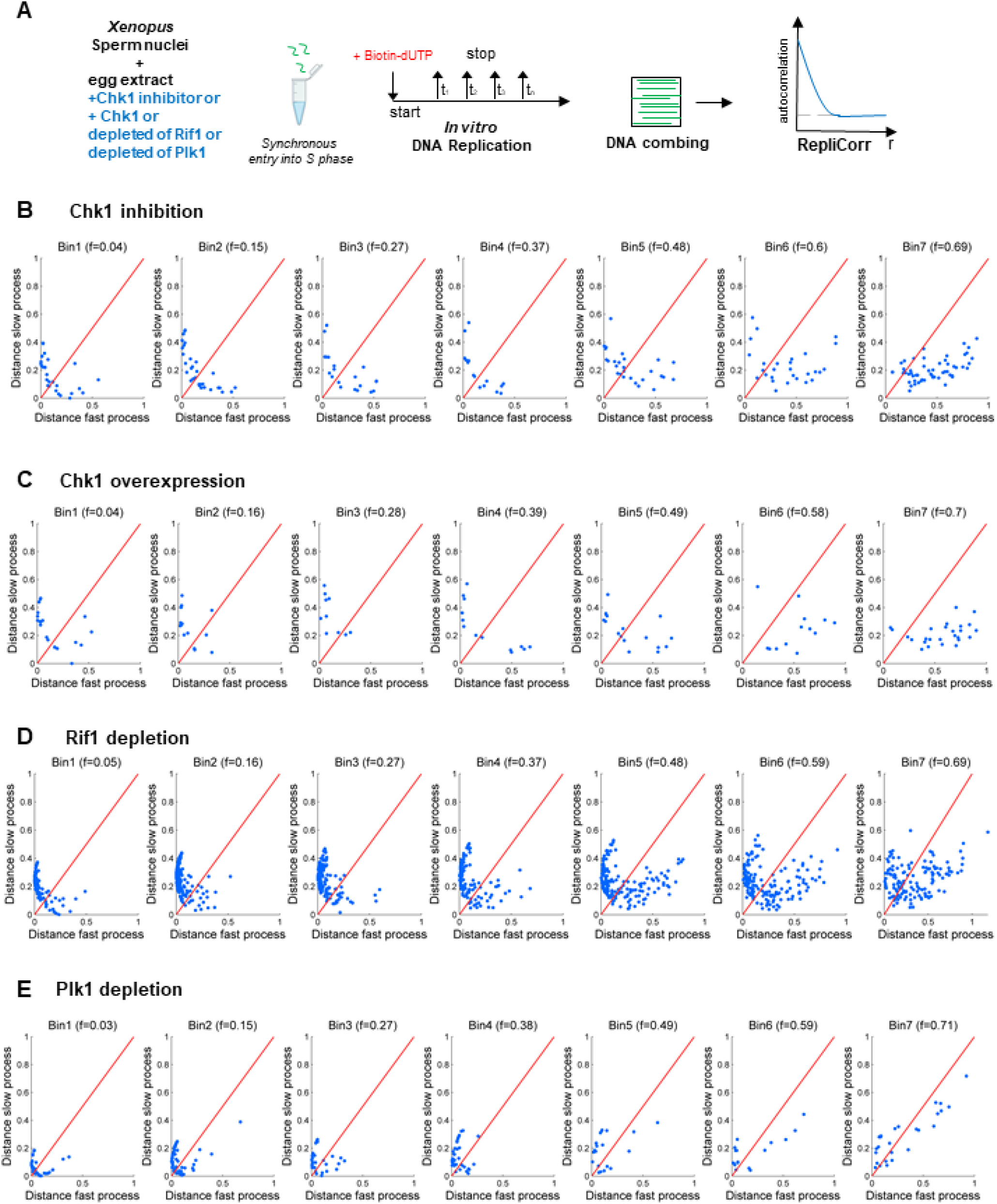
Modification of the replication patterns after Plk1 depletion but not after Chk1 inhibition, overexpression, or Rif1 depletion. (**A**) An outline of experimental workflow: sperm nuclei were incubated in Chk1 inhibited or Chk1 overexpressed egg extracts or Rif1 or Plk1 immunodepleted egg extracts in the presence of biotin dUTP. Genomic DNA was isolated at different times during the S phase, subjected to combing analysis, and further analyzed by RepliCorr. Normalized correlation coefficients (*ρ*_1_, *ρ*_2_) between the fiber’s *C*(*r, f*) and *C*1(*r, f*) and *C*2(*r, f*) were calculated. The similarity distance between the fiber and each process was defined as 1-*ρ*_1_ for the fast process and 1-*ρ*_2_ for the slow process and represented on a two orthogonal axis plot. The blue diagonal represents points of equal similarity to the two processes. (**B**) Chk1 inhibition by UCN-01 (independent experiments n=2). (**C**) Chk1 overexpression (n=2). (**D**) Rif1 depletion (n=2) and (**E**) Plk1 depletion (n=3).

## Discussion

In this study, we have explored how the activation of replication origins is coordinated along the chromosomes in a vertebrate model system. To address this question, we first developed a novel analysis method describing the spatial replication pattern of stretched single DNA molecules obtained by DNA combing or optical mapping after replication in the *Xenopus in vitro* system. We classified the similarity of these patterns, taking advantage of the correlation concept, and called this analysis method ”RepliCorr”. Second, our results reveal that two independent, spatio-temporally exclusive processes regulate DNA replication in *Xenopus*. These processes differ by their replication fork speed and rate of origin firing. Third, the abrogation of two main regulatory pathways of the DNA replication program, the replication checkpoint and Rif1 had either no or only a moderate influence on the spatial distribution of these processes. However, the depletion of the Polo-like kinase 1, known as checkpoint adaptor, abolished the spatial separation of these processes. Thus, our results suggest that Plk1 is an important coordinator of the spatial replication program and the initiation-elongation coupling along the chromosomes in *Xenopus*.

### The replication dynamics can be described as a combination of only two independent processes with distinct fork speeds and initiation rates in Xenopus

To analyze the dynamics of the stochastic replication process, detecting replicated tracks on individual DNA molecules allowed us to measure the time-dependent rate of DNA replication. In past studies, only initiation rates have been used to explore the replication process quantitatively. We previously reported that the initiation rate follows a typical bell-shaped curve during the S phase in several model organisms [8, 9, 10]. Since fork speeds are not necessarily constant throughout the S phase [1, 15, 5], we have now investigated how initiation and elongation are quantitatively connected to ensure S phase completion. Using RepliCorr, we show that the replication profiles of single DNA molecules during the normal S phase in *Xenopus* can be described by either of two processes, specified by the inverse relationship between initiation rate and fork speed. This confirms that initiation rate and fork speed are intimately linked properties of undisturbed DNA replication, as observed in mammalian cells [15]. Unexpectedly, we further demonstrate that the observed replication patterns can be described by a linear combination of two extreme configurations: one with a low initiation rate coupled with a fast fork progression and one with a high initiation rate associated with a slow fork progression (Figure 6A). We observe these two replication modes at the level of DNA molecules with 80-150 kb of size, corresponding to the size of replication clusters previously described in this experimental system [3, 5]. Therefore, the two replication modes may characterize two replication cluster types whose possible differences in chromatin structure or looping would be interesting to investigate. Are these two different replication strategies correlated with the temporal program? We observed that both processes co-exist during the early S phase, whereas the slow process becomes predominant as the S phase progresses. The change from a low to a higher initiation rate could be explained by initially limiting initiation factors, which, as the S phase progresses, are recycled towards origins to be activated in the unreplicated fraction of the genome. Fork speed could slow down during the S phase because of the progressive exhaustion of dNTPs at the nuclei concentration we used in the *in vitro* system and a low ribonucleotide reductase (RNR) activity expression. In early *Drosophila* embryos, the maternally deposited RNR is activated as dATP concentration decreases during the S phase [57]. However, decreasing fork speed in *Xenopus* was also observed at a ten times lower nuclei concentration [5], arguing against this explanation. Slow or stalled replication forks are considered as a sign of replication stress, which may result from DNA damage. Still, our observations suggest that impairment of the DNA replication checkpoint does not affect dual DNA replication modes. Finally, chromatin assembly can also regulate fork speed [58]. It is possible that the chromatin remodeling of sperm nuclei introduced into egg extracts creates a heterogeneous chromatin with co-existing accessible and difficult-to-replicate regions without activation of checkpoint mechanisms during DNA replication.

**Figure 6.**
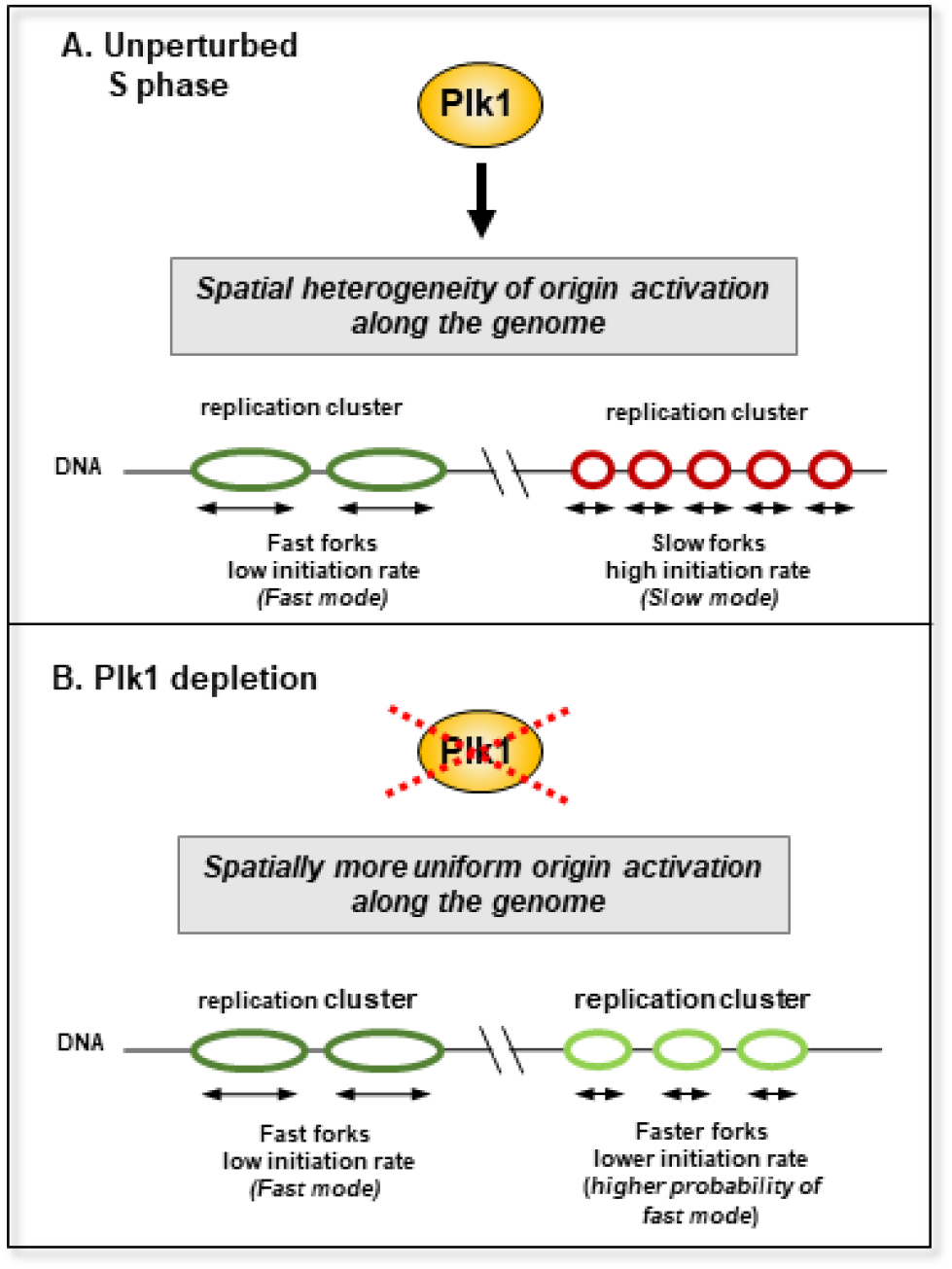
Model of a Plk1-dependent regulation of the spatial replication program by a fast and slow replication process in *Xenopus*. (**A**) In the presence of Plk1, two different replication patterns on DNA molecules can be distinguished, characterized by different fork speeds and initiation rates, leading to a non-uniform pattern of origin activation. (**B**) Upon Plk1 depletion, origin activation along the genome becomes more homogeneous; the slow replication mode approaches the fast mode.

### Polo-like kinase 1 regulates the spatial replication program

RepliCorr analysis of single-molecule replication patterns after targeting three different pathways of the replication program revealed that only Plk1 depletion strongly affected the pattern distributions between the fast and slow process. Without Plk1, these two processes were no longer independent, but they coexisted on DNA molecules with a high degree of replication, suggesting that Plk1 depletion canceled out the spatiotemporal exclusive character of the two processes (Figure 6B). In contrast, Rif1 depletion only moderately modified the pattern distribution, while Chk1 inhibition nor overexpression had any effect on this distribution. This is consistent with the fact that Plk1 depletion induced significant changes in initiation rates, IODs, and eye lengths when molecules of the same replicated fraction were compared [47], whereas Rif1 depletion [38] or Chk1 inhibition [31] did not. Therefore, these findings suggest that Plk1 promotes origin activation inside replication clusters. In contrast, Rif1 has been shown to mainly accelerate both whole cluster activation and replication of larger replication domains in *Xenopus* [38] and in mammalian cells [36, 37] consistent with only a small effect of Rif1 depletion on RepliCorr patterns detected in this study. Interestingly, our results suggest that Plk1 exclusively supports the slow process (Supplementary Figure S6 A) that predominates during the late S phase. Therefore, Plk1 may favor the late origin firing, similar to the dispersed, late firing origins identified in other eukaryotes [59, 60, 61, 62]. Chk1 inhibition or Rif1 depletion increases the initiation frequency of the slow process, albeit to a much lesser extent than Plk1 (Supplementary Figure S6 B, D), suggesting that the spatial regulation of the replication program by these known regulators mainly occurs via the regulation of the slow process in addition to their effects on the temporal program [47, 38].

It is unclear what could be the molecular mechanisms of how Plk1 locally regulates both fork speed and initiation rate in some genomic regions but not in others. Recently, we demonstrated that Plk1 could inhibit the Chk1-dependent replication checkpoint [46] and could phosphorylate the PP1 binding site of Rif1, which prevented PP1 inhibition by Rif1 in *Xenopus* [47]. However, we did not observe any effect on replication pattern distribution after Chk1 inhibition, and only a modest effect was seen after Rif1 depletion. Therefore, other Plk1-dependent pathways seem to be necessary for this local regulation. Interestingly, Plk1 co-immunoprecipitated with initiation complex proteins Treslin, MTBP, and TopBP1 [47], which are rate limiting for replication in *Xenopus* and budding yeast [63, 64]. In addition, Plk1 depletion results in a longer persistence of these factors on chromatin, and it has been suggested that these factors should dissociate from activated origins to allow fork elongation (conversion from pre-IC into CMG-complex) [65, 66]. It is, therefore, tempting to speculate that in the absence of Plk1, the recycling of Treslin/MTBP and TopBP1 towards neighboring origins is slowed down, resulting in a decrease in the initiation rate. In further support of this hypothesis, a recent study showed that for dormant origin firing, the linear correlation between IODs and fork speed at different concentrations of aphidicolin is also dependent on TopBP1 but not on Chk1 [67]. Another possibility is that Plk1 directly interacts with replication fork proteins to reduce fork speed. In favor of this possibility, we have shown that Plk1 co-immunoprecipitates with Rfc2-5, which is necessary to load the elongation factor PCNA. During the very early stages of *Xenopus* development, Plk1 levels are high but decline after the onset of zygotic transcription after the mid-blastula transition (MBT) [46]. The number of active origins also declined after the MBT [68, 69], but fork speed was not determined. During early mice developmental stages, mean origin distances gradually increase after the 2-cell embryo stage together with mean fork speed [24]. It would be interesting to investigate whether Plk1 could also be implicated in changing the dual replication modes during development.

In conclusion, our work shows that Plk1 promotes the spatio-temporal heterogeneity of initiation rate and fork speed. Plk1 is often over-expressed in aggressive cancer types [70] but is mainly studied for its role in mitosis entry. We believe it is necessary to consider the role of Plk1 during the S phase more carefully in tumor development. RepliCorr is a powerful tool for analyzing replication dynamics and initiating rate and fork speed coupling. It is robust enough to handle fluctuations resulting from the stochasticity of the replication process. The bell shape of the initiation rate was first observed in *Xenopus* [8] and later found to be universal in all eukaryotes [9, 10]. It would be of great interest to use RepliCorr analysis in single-molecule data from other model systems to see whether the basic principles observed in the *Xenopus* embryonic model can also apply to differentiated cells.

## Data availability

The DNA combing datasets analyzed during the current study are available from the corresponding author on request. The code for the RepliCorr analysis is deposited on GitHub (https://github.com/DidiCi/RepliCorr).

## Supplementary Data

Supplementary Methods, Supplementary Figures S1-6 and Tables S1-3 are available at NAR online.

## Competing interests

No competing interest is declared.

## Author contributions statement

K.M., O.H. (Saclay), and O.H. (Paris) conceived the experiment(s), C.D., O.H. (Saclay), F.dC. and K.M. conducted the experiment(s), and C.D., O.H. (Paris), K.M., and A.G. analyzed the results. K.M. wrote the manuscript with the input of the other co-authors.

## Acknowledgments

This work was supported by the IdEX Program of Paris-Saclay University for the interdisciplinary PhD fellowship (IDI) of D.C. to K.M., the Centre Nationale de Recherche Scientifique (CNRS, K.M., O.H., D.C.); the Commissariat a` l’Energie Atomique (CEA, A.G.), and in part by the Fondation de la Recherche Medicale (FRM, DEI20151234404, O.H., A.G.), Institut National du Cancer (INCa, PLBIO16-302, O.H., A.G.). and the Agence Nationale de la Recherche (ANR-15-CE12-0011-01, O.H.). We thank Mathis Miroux for technical assistance in LaTeX.

## Supplementary Data

-Supplementary Methods

-Supplementary Figures S1-6 and Tables S1-3

### Supplementary Methods

#### The theoretical framework

##### Stable phase fraction

The quantitative relation between the density of nucleated domains, the speed of growth, and the transformed volume during a nucleation and growth process were obtained between the end of the 30s and the beginning of the 40s by Kolmogorov, Johnson, Mehl and Avrami. The theory was pushed further by Sekimoto for the one-dimensional case [11]. In the KJMA model, the system undergoes a gradual transformation from an initial phase (metastable phase) to a final phase (stable phase), and the two phases coexist during the entire transition. In our case, the two phases correspond to the unreplicated and replicated state. During the transformation, stable domains nucleate randomly and grow in the metastable phase. The critical nucleus size, above which nuclei grow but below which they dissolve, is considered in the replication process to be infinitesimal. The process is characterized by *I*(*t*), the rate of nucleation per unit volume of metastable material, and 2*v*, the constant positive speed at which the stable phase grows after nucleation. We introduce the phase indicator function *u*(*r, t*), defined as follows:

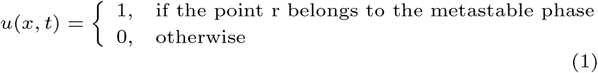

The fraction *ϕ*(*t*) for the metastable phase is then defined as:

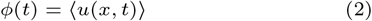

where ⟨⟩ denotes the average over the ensemble of the random variable *u*(*x, t*). *ϕ*(*t*) should be a decreasing function of *t*.

In order to obtain the formula for the metastable phase fraction, we must introduce the notion of a causal cone. This notion allows us to keep track of the complete history of the nucleation process, which is necessary in the case of continuous nucleation. The growth of the domain with constant speed 2*v*, from a specific nucleation site, can be viewed as an expanding triangle in the space-time representation. For multiple nucleations, the growth can be represented as the combination of different triangles, as shown in Fig. 1.

**Figure 1.**
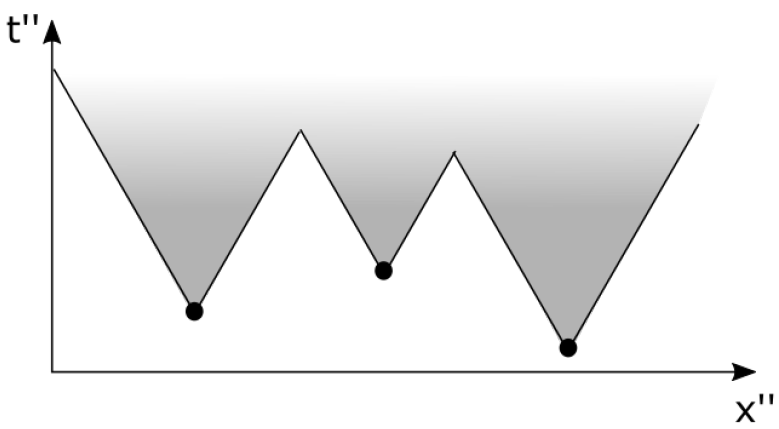
Nucleation and growth in one dimension. The stable domains that grow from multiple nucleation sites are unions of triangles in the spacetime representation; the resulting region is highlighted in grey.

On the other hand, for a point *x* to remain in the metastable phase at time *t*, nucleation events cannot occur within the inverted triangle, whose apex is at the point (*x, t*), as shown in Fig. 2. The inverted triangle is called the causal cone. As *ϕ*(*t*) corresponds to the probability that for any *t*^*′*^ *< t* nucleation centers do not appear in the length *S*(*t* − *t*^*′*^) = 2*v*(*t* − *t*^*′*^). Assuming nucleation as a rare event with a density *I*(*t*), we use the Poisson distribution to write:

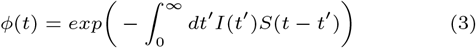

**Figure 2.**
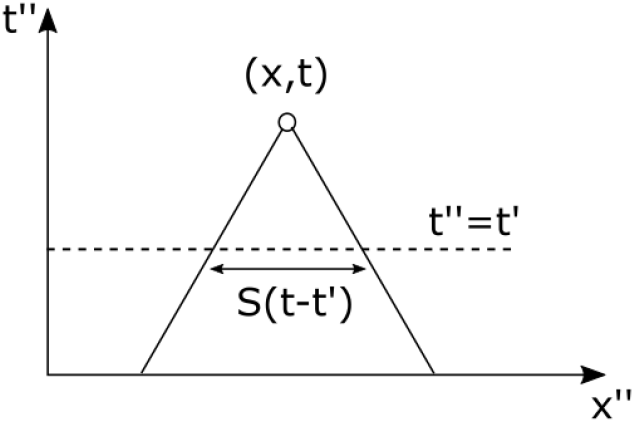
Representation of the one dimensional causal cone for the point (*x, t*).

with *S*(*t* − *t*^*′*^) = 0 for *t < t*^*′*^. This expression and what follows are valid only if: i) the speed *v* is not an increasing function of *t*, ii) the rate of nucleation is spatially homogeneous, and iii) the nucleation events occur independently. The fraction *f* (*t*) for the stable phase is:

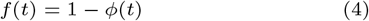

To have a more complete description of the nucleation and growth process, it is possible to analyze other quantities such as the length of islands, holes, and the island-to-island distances. The probability distribution of these quantities can be expressed as a function of the time t or the fraction of the stable phase f. In his work, Sekimoto also studied the time evolution of domain statistics by solving Fokker-Plank-type equations for island and hole distributions in the case of a constant nucleation rate I(t)=const [9, 10]. Sekimoto’s approach was extended in [6] in the case of a general nucleation rate I(t).

##### The correlation function

A further development of the theory was achieved by Sekimoto [11], who derived an exact expression for the two-point correlation function of growing domains in different dimensions. The correlation function provides in fact a more complete characterization of the spatial distribution of the two phases. Otha et al. [8] extended this result, introducing the possibility of nucleation of *p* different stable phases, and the limit of *p* → ∞ was analyzed by Axe and Yamada in one and two dimensions [4].

We derive the two-point correlation function in one dimension as deduced by Sekimoto [11] and Ohta, Ohta and Kawasaki [8]. The two-point correlation function can be expressed as:

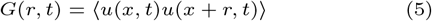

which quantifies the probability that two points separated by a distance *r* are both in the metastable phase at time *t*. In this case, the probability is governed by the union of the causal cones relatives to the two points (Fig. 3 and Fig. 4).

**Figure 3.**
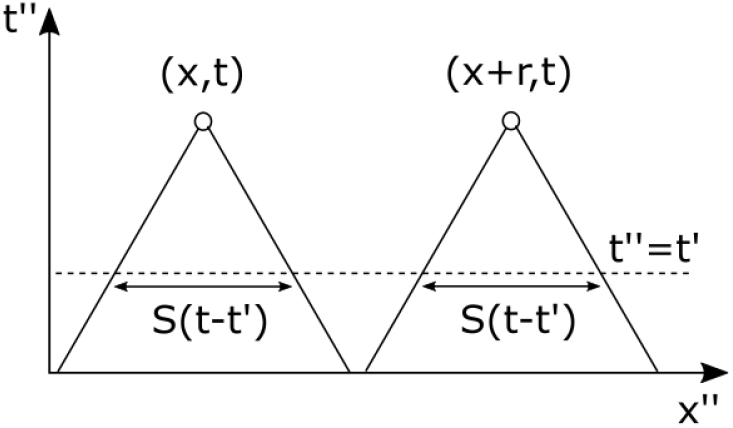
Representation of the causal cones for two uncorrelated points.

**Figure 4.**
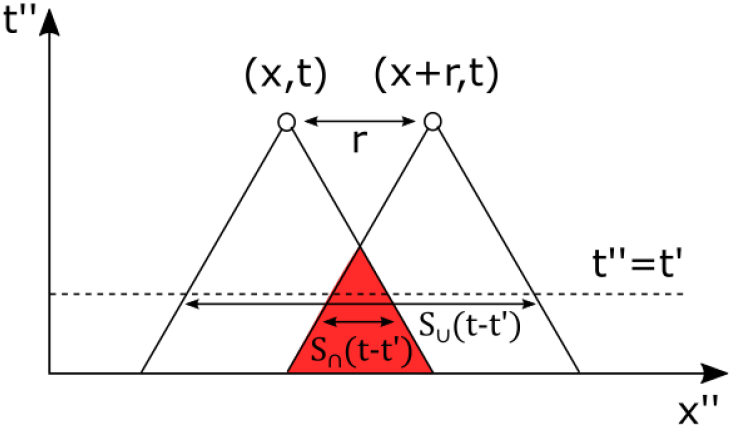
Representation of the causal cones for two correlated points. The region of overlap (in red) defines the degree of correlation between the two points.

With the same argument that we used for the evaluation of *ϕ*(*t*), we can write:

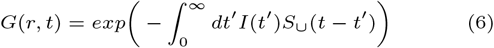

where *S*_∪_(*t* − *t*^*′*^) is the spatial length, in which no nucleation events must occur at time *t*^*′*^ *< t* in order that the points *x* and *x* + *r* belong to the metastable phase at time *t*. We have:

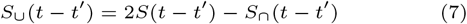

where *S*_∩_(*t* − *t*^*′*^) = 2*v*(*t* − *t*^*′*^) − *r* is the spatial length relative to the eventual intersection of the two causal cones and is equal to zero for *r >* 2*v*(*t* − *t*^*′*^). By substituting (7) in (6), we obtain:

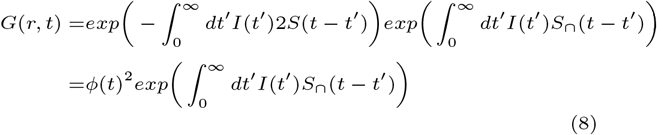

For *r >* 2*vt, G*(*r, t*) = *ϕ*(*t*)^2^ meaning that the phase state of two points at distance *r >* 2*vt* are independent. This reflects the fact that at the time *t*, the maximum size of a stable domain is 2*vt* and no long-range correlation is mediated by the stable domains.

We can then easily obtain the two-point correlation function for the stable phase as:

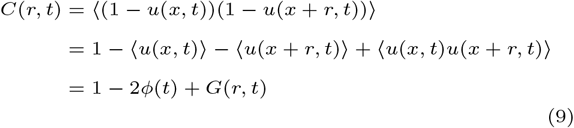

##### Correlation function of fluorescence profiles from DNA fiber experiments

As detailed above, Kolmogorov, Johnsol, Mehl and Avrami developed a stochastic model that describes the kinetics of the transition from an initial phase (metastable phase) to a final phase (stable phase). [7, 5, 1, 2, 3]. A further development of the theory was achieved by Sekimoto [11], who derived an exact expression for the two-point correlation function of growing domains in different dimensions. The KJMA theory can be used to describe the replication process if the replicated state is considered as the stable phase and the unreplicated phase as the metastable phase. In this context, the growth speed *v* corresponds to the replication fork speed, and the nucleation rate *I*(*t*) to the frequency of initiation. Once obtained an explicit form for the two-point correlation function, we applied it to the study of the correlation function of fluorescence intensity profiles from DNA fiber experiments.

In order to use the expressions (4) and (9), we need to choose an explicit form for the frequency of initiation *I*(*t*). We will consider the form:

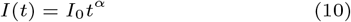

with *I*_0_ ≥ 0 and *α* ≥ 0. This expression is a good approximation for the increasing region of the frequency of initiation. We then restricted the analysis to this region. By using the Eq. (3) and (4), we obtain:

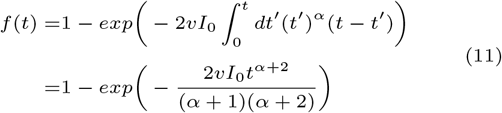

where we used *S*(*t* − *t*^*′*^) = 0 for *t < t*^*′*^. In a similar way, from the Eq. (8) and (9), we have:

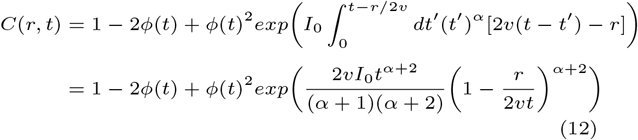

where we used *S*_∩_(*t*−*t*^*′*^) = 0 for *r >* 2*v*(*t*−*t*^*′*^) or equivalently for 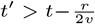. The Eq. (12) will be valid for *r < l*_*max*_ = 2*vt*, where *l*_*max*_ represents the maximum replication eye length present at time *t*.

##### Statement of the problem

The DNA combing and HOMARD technique allows the analysis of the replication state of a DNA fiber at a certain time during the replicative phase. Between all the information that we can obtain through the analysis of the fluorescence intensity profiles, we will focus on the replicated fraction, the frequency of initiation, and the correlation function of the single fiber. In the analysis, there will be two major consequences related to the use of data obtained with the DNA combing technique:

1. The finite size of the analyzed fibers implies that we only have access to the local evolution of the process. The results of the quantitative analysis will be not valid for the replication process of the entire genome.
2. To apply the theory as it is, we would need to know the exact time at which replication starts on the single fiber, but the experimental technique does not provide this information. We will have to use the replicated fraction as a measure of the local evolution of the process. This will introduce some uncertainty in the analysis, due to the lack of knowledge of the hidden variable, that is, the time.

We can define the problem as follows. We want to analyze the similarity between replication patterns of different fibers by comparing the correlation function of the fluorescence intensity profiles. The specific pattern depends on the frequency of initiation and the fork speed. So, we will estimate the variables *v, I*_0_ and *α*, given the experimental frequency of initiation *I*(*f*) as a function of the replicated fraction *f* and the correlation function *C*(*r, f*) for different replicated fractions *f* as a function of *r*. The time will be obtained from the analytical inversion of the Eq. (11) as *t* = *f*^*−*1^(*v, I*_0_, *α*).

## Supplementary Figures and Tables

**Supplementary Table S1:** Comparison of fit results for correlation function with one process between simulated and experimental data. The rate of initiation is I(t)=I_0_t_***α***_ per unit time per length of unreplicated DNA. Therefore, the replication process is characterized by three parameters: *v*, and for the initiation rate, *I*_*0*_, ***α*·** Results of fit with a constant *I*_*0*,_ ***α*** and fork speed *v* for the autocorrelation function of simulated data from Figure 1C and from fit to experimental data from Figure 1D. The parameter values were averaged over 100 trials. A χ_2_ value close to 1 is considered as a very good fit.

| | $v$<br>(kb/min) | $l_0$<br>(1/(kb*min<sup>$\alpha+1$</sup>)) | $\alpha$ | $\chi^2$ |
| --- | --- | --- | --- | --- |
| Simulation parameters | 1 | 0.03 | 0 |  |
| Fit parameters |  |  |  |  |
| Simulated replication data Fig. 1C | 1.009 | 0.029 | 0 | 2.2 |
| Fit parameters |  |  |  |  |
| Experimental data Fig. 1D | 0.76 | 0.000029 | 1.98 | 13.1 |

**Supplementary Figure S1:**
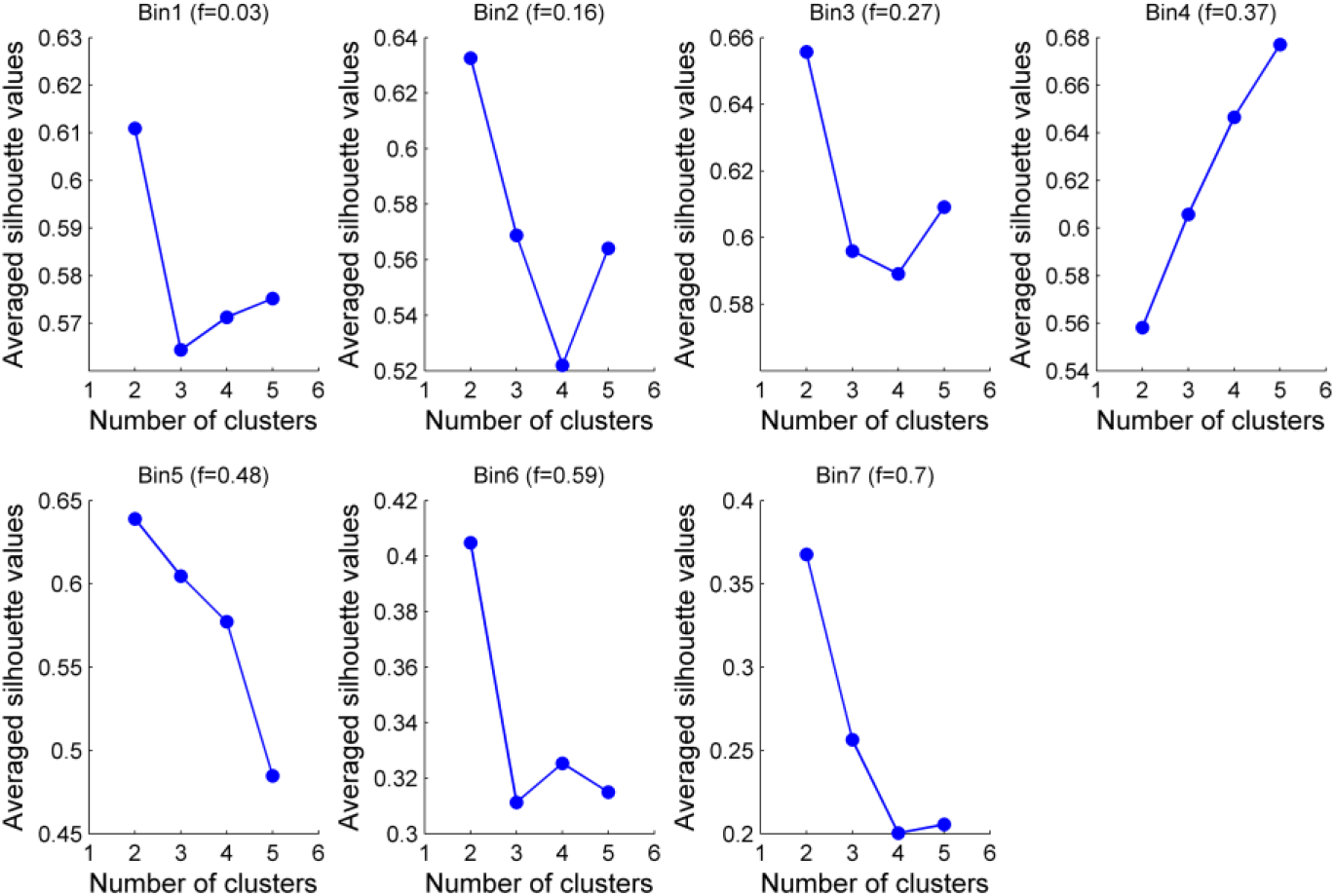
Clustering evaluation by the average of silhouette values for the first control set. Each image corresponds to a different interval of replicated fractions from 0 to 75% (Bin 1-7) and the averaged replicated fraction is reported on the top. The fibers in each interval of replicated fraction were grouped into two to five clusters and the silhouette value was calculated for each configuration.

**Supplementary Figure S2:**
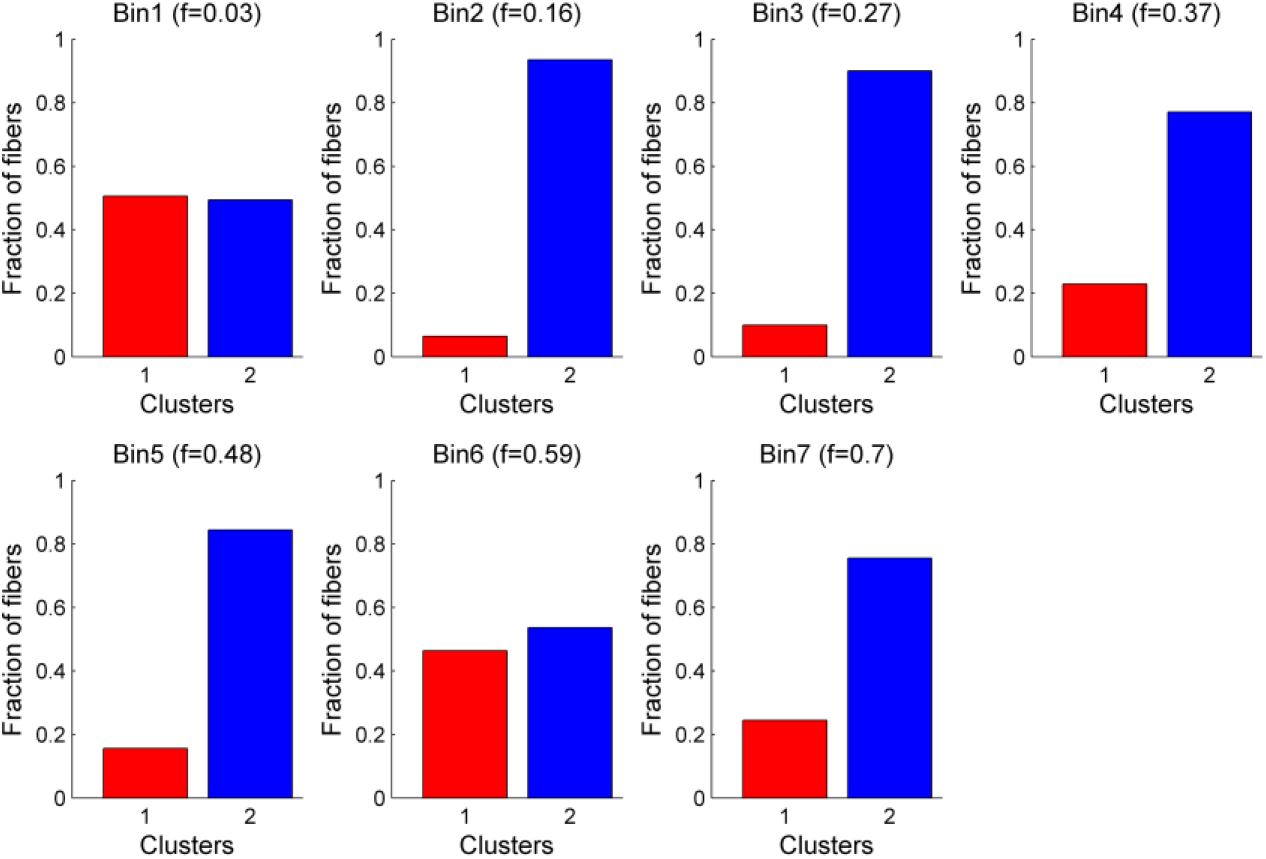
Histogram reporting the number of fibers in each cluster for the first control set. Each image corresponds to a different interval of replicated fractions from 0 to 75% (Bin 1-7) and the averaged replicated fraction is reported on the top.

**Supplementary Table S2:**
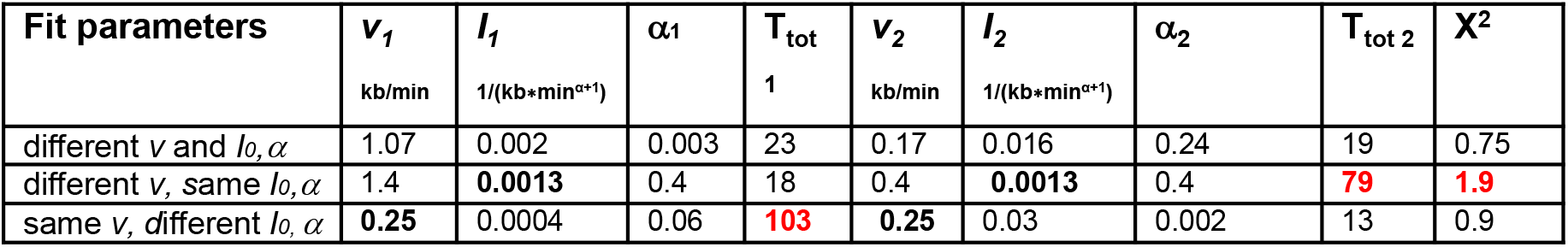
Fitting parameters values from Figures 3A with 2 processes with varying *v* and *I0* compared to fitting parameters with constant *I*_*0*_, ***α***, or constant *v* from Supplementary Figure S4 A and B. The initiation rate is given by I(t)=I_0_*t_***α***_ per unit time per length of unreplicated DNA. Therefore, each process is characterized by three parameters: *v, I*_*0*_, ***α***. For different I(t) values of I_0_ and ***α*** are set or parameters given after fitting. *I*_*1*_ and *I*_*2*_ are *I*_*0*_ for process 1 and 2, respectively. T_tot_ is the time in min to replicate a fiber to 75 %. Values in red are those differing from the fit with two different *I*_*0*,_ ***α*** and *v*.

**Supplementary Figure S3:**
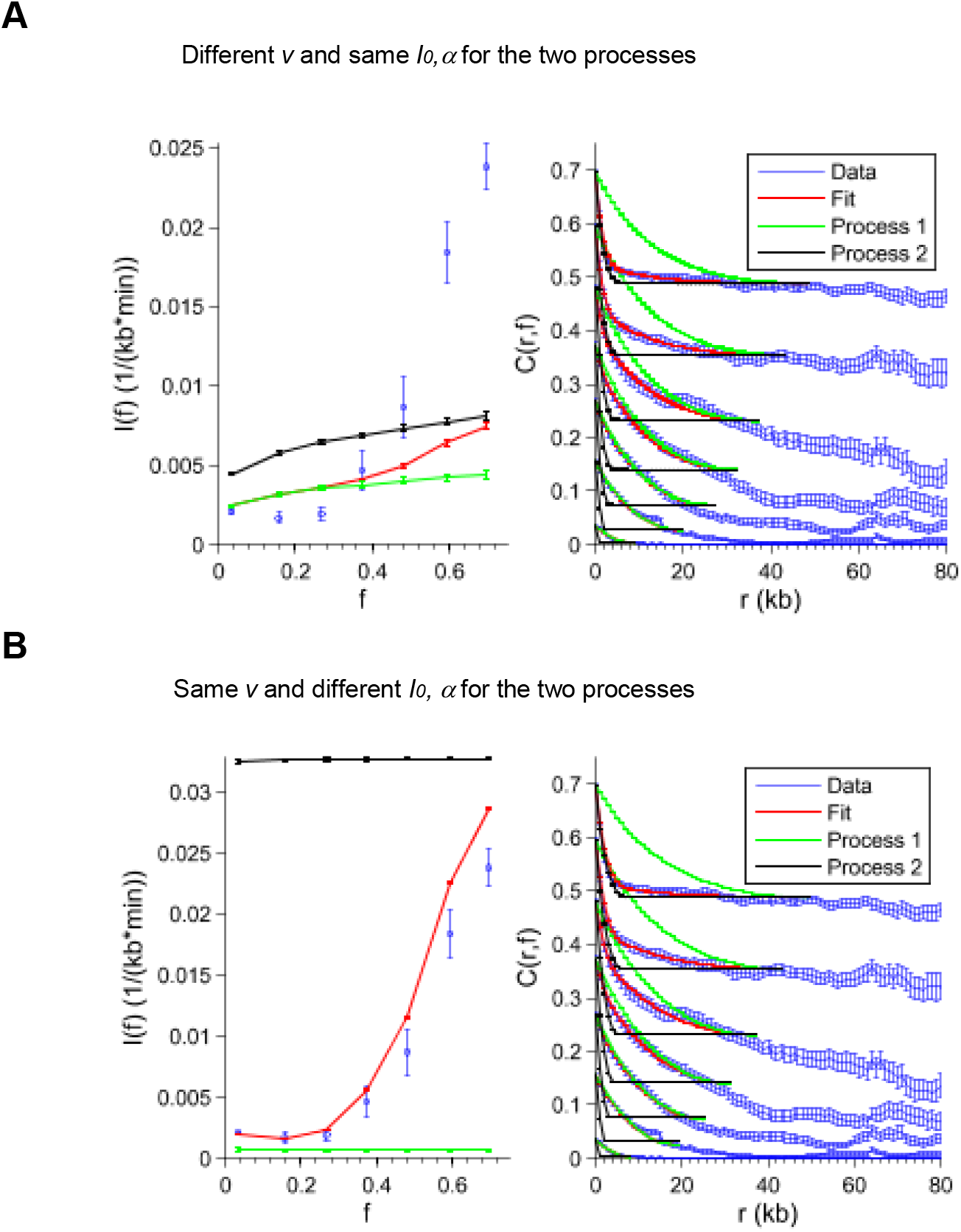
Comparison between fits with two processes for correlation profiles with different or same *I*_*0*,_*α* and *v*. (**A**) with different *v* and same *I*_*0*_. **(B**) with same *v* and different *I*_*0*_ for the two processes. (**C**) table with fitted parameter values from A and B compared to the parameters from Figure 3 A; with T_tot_ being the time in min to replicate a fiber to 75 %. We highlight values in red those differing from the fit with two different *I*_*0*_ and *v*. The parameter values of fork speed and initiation frequency were averaged over 100 trials.

**Supplementary Figure S4:**
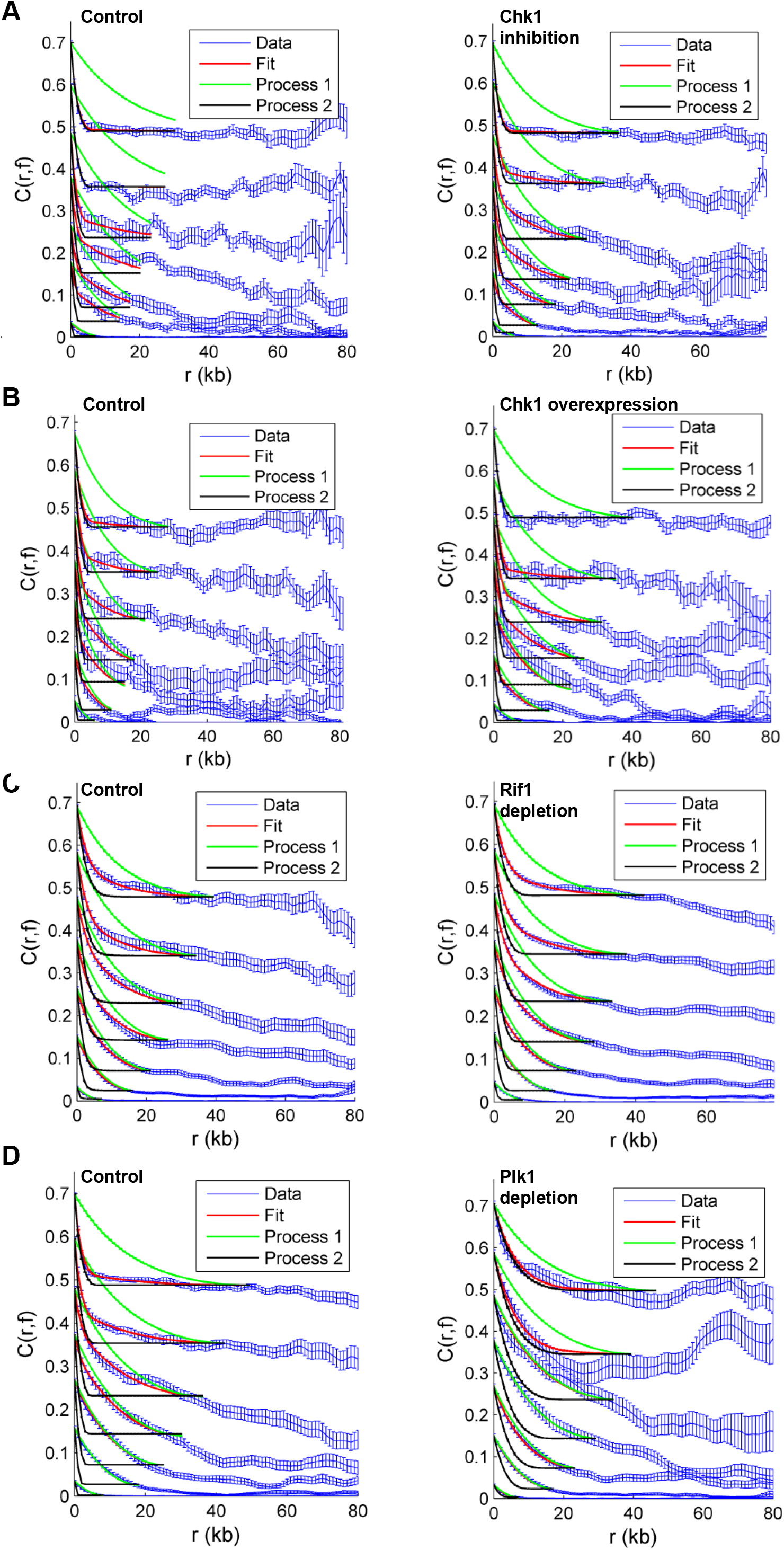
Mean autocorrelation function *C(r*,*f)* profiles (blue curve, with standard deviation) with fit (red curve) as in Figure 3A for different pathways perturbations and corresponding controls. The green curve is the correlation profile produced by the fast fork process (*C1(r*,*f)*, model 1), and the black curve is the correlation profile produced by the slow fork process (*C2(r*,*f)*, model 2), f. (**A**) Chk1 inhibition by UCN-01 (n=2). (**B**) Chk1 overexpression (n=2). **(C)** Rif1 depletion (n=2). **(D)** Plk1 depletion (n=3).

**Supplementary Figure S5:**
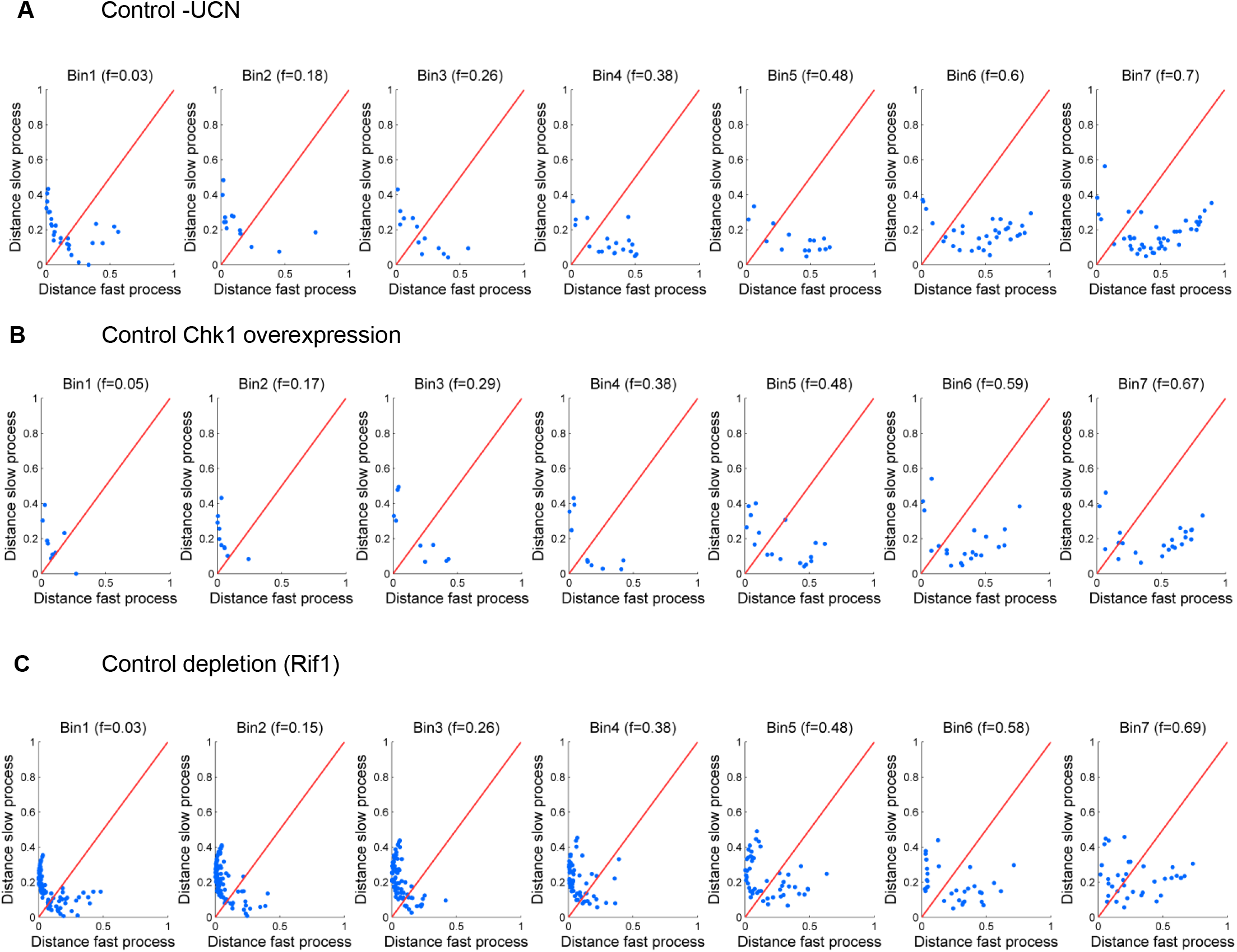
Similarity distances of the fast and the slow process from control experiments from Figure 5. Normalized correlation coefficients (ϱ1, ϱ2) between the fiber’s *C(r*,*f)* and *C*_*1*_*(r*,*f)* and *C*_*2*_*(r*,*f)* were calculated for control conditions in Figure 5 B-D. The similarity distance between the fiber and each process was defined as 1-ϱ1 for fast process and 1-ϱ2 for slow process and represented on a two orthogonal axis plot. The red diagonal represents points of equal similarity to the two processes. (**A**) +DMSO as control condition for Chk1 inhibition by UCN. (**B)** Control for Chk1 overexpression protein buffer addition. (**C**) Control depletion for Rif1 experiment.

**Supplementary Figure S6:**
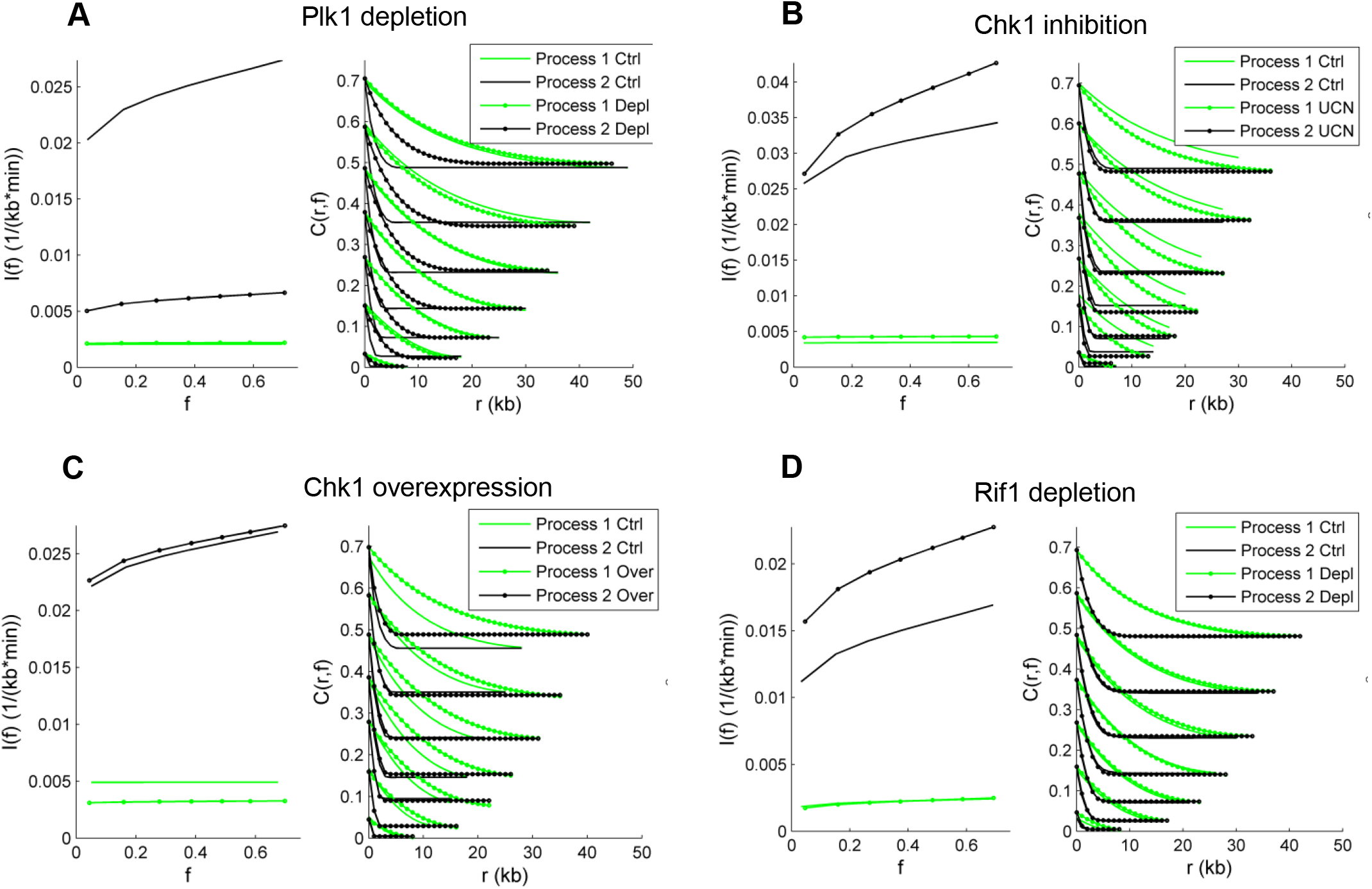
Fits for correlation functions *C(r*,*f)* and associated initiation rates for fast process (process 1, green curves) and slow process (process 2, black curves) from experiments of (**A**) Plk1 depletion and its control. (**B**) UCN inhibition and control. (**C**) Chk1 overexpression and control. (**D**) Rif1 depletion and control.

**Supplementary Table S3:** Values from fits to autocorrelation profiles with two processes (1= fast process, 2=slow process) under different experimental conditions. The rate of initiation is I(t)=I_0_t_***α***_ per unit time per length of unreplicated DNA. Therefore, the replication process is characterized by three parameters: *v*, and for the initiation rate, *I*_*0*_, ***α*·** I_1_ and I_2_ are the I_0’s_ for each process. The parameter values of fork speed and initiation frequency were averaged over 100 trials.

|  | Fast process |  |  | Slow process |  |  |  |
| --- | --- | --- | --- | --- | --- | --- | --- |
| | $v_1$ | $l_1$ | $\alpha_1$ | $v_2$ | $l_2$ | $\alpha_2$ | $\chi^2$ |
| Experimental conditions | (kb/min) | 1/(kb*min<sup>$\alpha+1$</sup>) |  | (kb/min) | 1/(kb*min<sup>$\alpha+1$</sup>) |  |  |
| Control -UCN | 2.251 | 0.003 | 0.009 | 0.211 | 0.021 | 0.197 | 1.14 |
| Chk1 inhibition + UCN | 1.556 | 0.004 | 0.014 | 0.191 | 0.020 | 0.356 | 0.40 |
| Control Chk1 overexpression | 1.141 | 0.005 | 0.002 | 0.138 | 0.018 | 0.178 | 0.63 |
| +Chk1 | 1.333 | 0.003 | 0.032 | 0.167 | 0.019 | 0.155 | 0.68 |
| Control depletion (Rif1) | 0.807 | 0.002 | 0.160 | 0.208 | 0.008 | 0.383 | 0.86 |
| Rif1 depletion | 0.767 | 0.001 | 0.160 | 0.297 | 0.011 | 0.303 | 1.05 |
| Control depletion (Plk1) | 1.073 | 0.002 | 0.003 | 0.170 | 0.016 | 0.236 | 0.74 |
| Plk1 depletion | 1.034 | 0.002 | 0.022 | 0.471 | 0.004 | 0.209 | 0.93 |

## Notes

### Competing Interest Statement

The authors have declared no competing interest.

